# Cryo-EM structural insights into Vγ9Vδ2 TCR activation via multiple butyrophilins

**DOI:** 10.1101/2024.10.02.616253

**Authors:** Mai Zhang, Yiqing Wang, Ningning Cai, Yingying Qu, Xianqiang Ma, Jing Xue, Xiaorui Chen, Xueguang Zhang, Junyu Xiao, Yonghui Zhang

**Author notes:** Corresponding author. (Y.Z.); (J.X.). These authors contributed equally to this work (Mai Zhang, Yiqing Wang, Ningning Cai, Yingying Qu).

## Abstract

Human Vγ9Vδ2 T cells detect cancerous and pathogen-infected cells through an “inside-out” mechanism wherein intracellular domains of butyrophilin (BTN) proteins recognize phosphoantigens (pAgs) to prime BTN ectodomains to engage with the T cell receptor (TCR) to activate Vγ9Vδ2 T cells. Here, we present cryo-electron microscopy (cryo-EM) structures, including the full-length BTN3A1-BTN3A2–BTN2A1 complex stabilized by the pAg HMBPP and an HMBPP-primed BTN multimer complex engaged with the Vγ9Vδ2 TCR. The HMBPP-primed BTN3A1-BTN3A2–BTN2A1 complex confirms that this pAg bridges the intracellular B30.2 domains of BTN3A1 and BTN2A1, while also revealing an association of the ectodomains of BTN3A2 and BTN2A1. Upon TCR engagement, the TCR undergoes a substantial conformational change and BTN3A2–BTN2A1 ectodomain interaction dissociates, allowing BTN2A1 to bind to the lateral surface of the Vγ9 chain, while BTN3A2 binds to the apical surface of the Vδ2 chain. Beyond demonstrating that BTN3A2 as a *bona fide* TCR ligand, this dissociation/binding sequence suggests a “pliers-like gripping” mechanism for TCR activation. We also obtained cryo-EM structures of BTN proteins in complex with functional antibodies, further elucidating the overall Vγ9Vδ2 T cell activation process. Thus, our structural insights into how BTN proteins sense antigens and drive TCR activation lay the groundwork for developing targeted γδ T cell immunotherapies.

## INTRODUCTION

T cells are broadly categorized into two types based on the structure of their T cell receptors (TCRs): αβ T cells and γδ T cells. αβ T cells recognize antigens that bind within a groove formed by the extracellular region of major histocompatibility complex (MHC) or MHC-like molecules, facilitating direct interactions with the αβ TCRs^1^. In contrast, γδ T cells operate independently of the MHC^2,3^. A prominent subset of human circulating γδ T cells, Vγ9Vδ2 T cells, are known respond to infections and tumors by recognizing target cells that contain activating “phosphoantigens” (pAgs). pAgs are small isoprenoid metabolites including the metabolically universal isoprenoid precursors isopentenyl diphosphate (IPP) and dimethylallyl diphosphate (DMAPP); there are also more potent pAgs like (*E*)-1-hydroxy-2-methyl-but-2-enyl 4-diphosphate (HMBPP), which is produced by exogenous sources including bacteria and parasites^4–8^.

γδ T cell receptors (TCRs) are known to recognize multiple butyrophilin (BTN) or butyrophilin-like (BTNL/Btnl) proteins on target cells^9^. BTNs are transmembrane proteins with extracellular domains that resemble immunoglobulin variable (IgV) and constant (IgC) regions, and often include an intracellular B30.2 domain^10^. Studies of Vγ9Vδ2 T cell activation have established that the expression of BTN3A1 on target cells is prerequisite for sensing pAgs^11^, and BTN2A1 is also understood as necessary for pAgs sensing by Vγ9Vδ2 T cells^12,13^. Moreover, studies focused on BTN3A1 have indicated that the additional presence BTN3A2 or BTN3A3 is required for pAgs sensing^9,14,15^.

Intracellularly, BTN3A1 binds pAgs^16,17^ and forms a composite interface that facilitates interaction with BTN2A1, inducing “inside-out” conformational change^15^. Extracellularly, BTN2A1 engages the lateral surface of the Vγ9Vδ2 TCR, leaving the apical surface of the TCR unoccupied^18^. Notably, epitope mapping studies have suggested the existence of a second ligand-binding region, distinct from the BTN2A1 binding site, which is essential for pAgs reactivity^12,19,20^. However, it remains unclear whether a BTN3A subfamily protein—or perhaps some other accessory molecule—functions as an additional TCR ligand^18^.

In this study, we present multiple cryo-electron microscopy (cryo-EM) structures, including the full-length BTN3A1-BTN3A2–BTN2A1 complex stabilized by HMBPP and an HMBPP-primed BTN3A1-BTN3A2–BTN2A1 complex engaged with the Vγ9Vδ2 TCR. Structural analysis of the “primed” BTN3A1-BTN3A2–BTN2A1 complex confirmed that HMBPP glues the intracellular B30.2 domains of BTN3A1 and BTN2A1. Combined structural and cellular activation assay results revealed that the ectodomains of BTN3A2 and BTN2A1 interact, doing so in a manner that conceals binding epitopes for the Vγ9Vδ2 TCR, with the BTN3A1–BTN2A1 ectodomain interaction persisting up until TCR engagement occurs. Our TCR-engaged complex structure revealed that TCR engagement drives dissociation of the BTN3A1 and BTN2A1 ectodomains and shows how the TCR undergoes a conformational change that is driven by interactions between its Vδ2/Vγ9 chains and BTN3A2. Beyond experimentally confirming that BTN3A2 is a *bona fide* ligand of the Vγ9Vδ2 TCR, our structure reveals a fascinating “pliers-like gripping” TCR activation mechanism through which heterogenous BTN proteins engage separately with the lateral and apical sides of the Vγ9Vδ2 TCR. Finally, we obtained two additional cryo-EM structures for complexes for a BTN3A agonist monoclonal antibody (TH001) and a BTN2A1 antagonistic monoclonal antibody (TH002) in complex with BTN proteins, and harnessed these to explore the Vγ9Vδ2 T cell activation process.

## RESULTS

### Cryo-EM structure of the HMBPP-primed BTN3A1-BTN3A2–BTN2A1 complex

We first expressed and purified full-length BTN2A1 homodimer along with the BTN3A1–BTN3A2 heterodimer from HEK293F cells. Notably, we were only able to obtain sufficient full-length BTN3A1 when it was co-expressed with BTN3A2, a finding that aligns with two previous observations: that BTN3A2 enhances BTN3A1 expression^9,14^ and that the absence of BTN3A2 impairs the responsiveness of Vγ9Vδ2 T cells to pAgs^9,14,15^. Additionally, we purified the ectodomain (ECD) of the Vγ9Vδ2 TCR (clone G115^21^). Briefly, immunoblotting of *in vitro* assay products conducted to examine complex assembly showed that no complex forms in the absence of the pAg molecule HMBPP. In contrast, the presence of HMBPP induces assembly of the BTN3A1-BTN3A2––BTN2A1 complex (henceforth termed the “HMBPP-primed BTN multimer”); and in the presence of both HMBPP and purified Vγ9Vδ2 TCR, a “TCR-engaged” BTN3A1-BTN3A2–BTN2A1 complex assembles (Extended Data Fig. 1a).

For cryo-EM analysis, we initially focused on the HMBPP-primed BTN multimer complex. The final reconstructions revealed the extracellular region at 4.17 Å resolution, as well as the transmembrane (TM), intracellular juxtamembrane (JM) and B30.2 regions at 3.58 Å resolution (Extended Data Fig. 2 and Extended Data Table 1). Notably, both the BTN2A1 homodimer and the BTN3A1–BTN3A2 heterodimer form V-shaped dimeric structures (Fig. 1a). Within the two dimers, BTN monomers interact with each other via IgC domains in the extracellular region, burying an area of 420 Å² in the BTN2A1 homodimer and of 820 Å² in the BTN3A1–BTN3A2 heterodimer (Extended Data Fig. 3). The TM and JM regions of the BTN3A1–BTN3A2 and BTN2A1 molecules create continuous coiled-coil structures of approximately 120 Å in length (Fig. 1b).

**Fig. 1.**
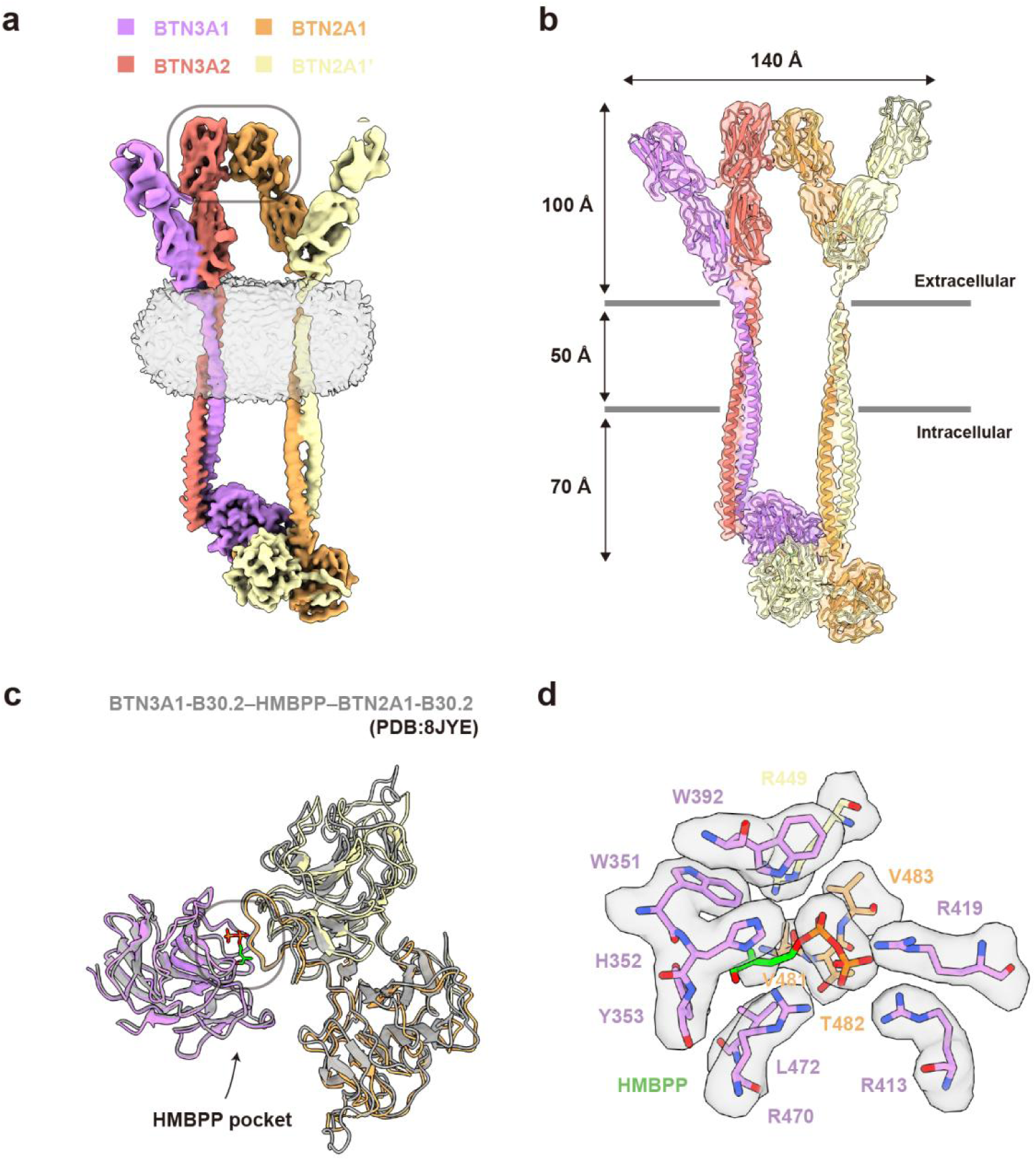
Cryo-EM structure of the HMBPP-primed BTN3A1-BTN3A2–BTN2A1 complex. **a,** Cryo-EM reconstruction of the complex. BTN3A1 and BTN3A2 are shown in purple and salmon, respectively; the two BTN2A1 molecules comprising the homodimer are shown in yellow and light yellow. The detergent micelles are depicted in transparent gray. The gray box indicates the IgV interface between BTN3A2 and BTN2A1. **b,** The TM/JM regions of the BTN3A1-BTN3A2 heterodimer and the BTN2A1 homodimer form coiled-coil structures. Models of BTN molecules and the cryo-EM reconstruction are shown. **c,** HMBPP facilitates the assembly of the BTN3A1-BTN3A2–BTN2A1 complex. The BTN3A1-B30.2–HMBPP–BTN2A1-B30.2 model resembles a previously determined crystal structure (PDB ID: 8JYE). **d,** The basic pocket that engages with HMBPP in the cryo-EM reconstruction. The pAg molecule HMBPP is shown as a green stick.

In the intracellular regions, the two BTN2A1 B30.2 domains constitute a homodimer and interact with one HMBPP-bound BTN3A1 B30.2 domain (Fig. 1c). BTN3A2 features a shorter cytosolic region and lacks the B30.2 domain. Consistent with our previous findings^15^, engagement between HMBPP and BTN3A1 B30.2 forms a composite interface for direct binding to BTN2A1 B30.2, with HMBPP centrally positioned at the interface. Specifically, the diphosphate group resides primarily in the basic pocket of BTN3A1 B30.2 and interacts with two BTN2A1 B30.2 domains (Fig. 1d), acting as a molecular glue to stabilize the complex.

### Interaction of the BTN3A and BTN2A1 ectodomains in the HMBPP-primed BTN3A1-BTN3A2–BTN2A1 complex

Previous studies have established that in the resting state, BTN2A1 and BTN3A1 are located closely (within 10 nm of each other) in *cis* at the cell surface, and the ectodomains of BTN3A and BTN2A1 engage in direct interaction, supporting a stable association^12,18^. A crystal structure of BTN3A1–BTN2A1 ectodomain heteromeric complex, tethered together with C-terminal leucine zippers, has been reported^18^.

Our structure of the HMBPP-primed BTN multimer reveals that the ectodomains of BTN2A1 and BTN3A2, but not BTN3A1, continue to interact upon pAg priming, specifically through the CFG faces of their IgV domains (Fig. 2a). This observation is supported by the previously reported preference of BTN3A2 to dimerize with BTN2A1 over BTN3A1^14,22^. The interface area is about 620 Å². And consistent with this interaction, a total of fourteen BTN3A2 residues are buried in the HMBPP-primed structure (Glu35, Ser41, Ser42, Arg44, Lys94, Leu96, Tyr98, Gln100, Gly102, Asp103, Tyr105, Glu106, Lys107, and Leu109) (Extended Data Fig. 4a). Similarly, seventeen BTN2A1 residues are buried (Glu35, Arg37, Phe39, Ser41, Gln42, Phe43, Ser44, Pro45, Lys51, Arg56, Glu59, Tyr98, Gln100, Gly102, Arg103, Tyr105, and Glu107) (Extended Data Fig. 4b).

**Fig. 2.**
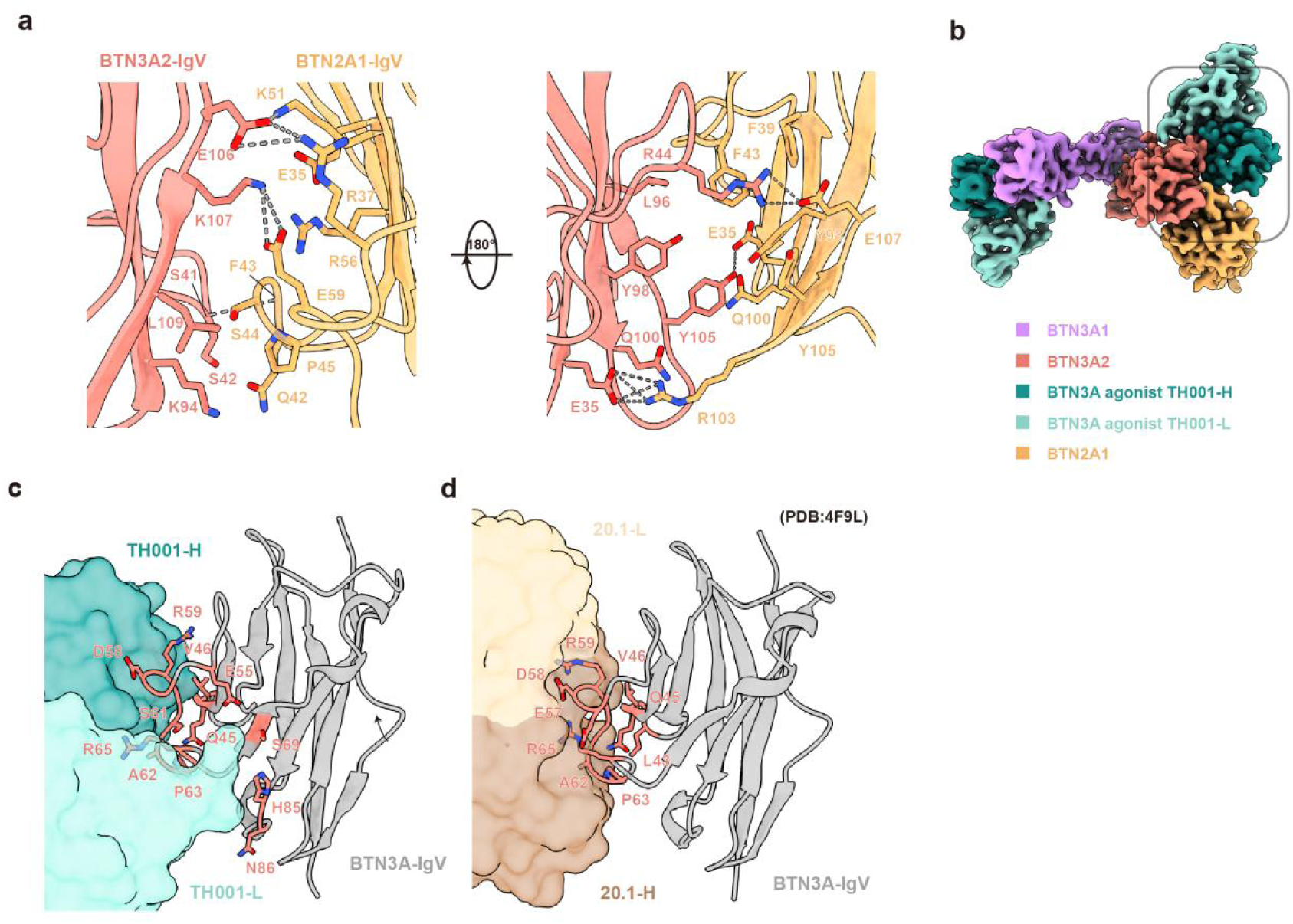
Interactions of the BTN3A and BTN2A1 ectodomains. **a,** Interactions between BTN3A2-IgV and BTN2A1-IgV in the HMBPP-primed BTN multimer. Dashed lines indicate polar interactions. **b,** Cryo-EM reconstruction of the BTN3A1-BTN3A2–BTN2A1–TH001-Fab complex. BTN3A1, BTN3A2, and BTN2A1 are shown in purple, salmon, and yellow, respectively. The heavy and light chains of TH001 are shown in two shades of green. The gray box indicates that TH001 has no effect on the interaction of BTN3A2–BTN2A1 ectodomains. **c-d,** Comparison of epitopes between two agonist BTN3A antibodies (TH001 and 20.1, the latter being a previously reported tool BTN3A agonistic antibody^30^. Antibodies are shown as surface representations. BTN3A molecules are shown as grey ribbons; residues of the epitopes are shown in salmon.

When considering in terms of a previously reported BTN2A1–Vγ9Vδ2 TCR structure (PDB ID: 8DFW), it is conspicuous that some of the buried BTN2A1 residues in our HMBPP-primed BTN multimer complex also participate in BTN2A1’s interaction with the Vγ9 TCR. This observed overlap in the BTN2A1 residues that engage with BTN3A2 and with Vγ9 TCR may clarify key concepts in the γδ T cell activation field, such as whether separation of the BTN3A1 and BTN2A1 ectodomains is essential for TCR activation, and/or how pAgs priming might result in the exposure of extracellular epitopes for Vγ9Vδ2 TCR engagement.

Our HMBPP-primed BTN multimer complex supports that, even in the presence of HMBPP, the interaction between the IgV domains of BTN3A1 and BTN2A1 conceals residues that are later involved in TCR engagement. Accordingly, it would follow that dissociation of the BTN3A–BTN2A1 ectodomains may only occur upon TCR engagement. Supporting this, a previous study reported that locking the ectodomains of BTN3A1 and BTN2A1 together abrogates their interaction with the Vγ9Vδ2 TCR^18^. Conversely, Vγ9Vδ2 T cell activation is significantly enhanced upon introducing mutations to BTN2A1 IgV domain that weaken its association with the BTN3A2 IgV domain, as evidenced by cellular assays conducted in the present study (E59A mutation) and our previously reported work^15^ (E35A and R56A) (Extended Data Fig. 4c). Collectively, our structural and assay results support that the interaction between the BTN3A1 and BTN2A1 ectodomains persists until engagement with the Vγ9Vδ2 TCR.

We next examined whether TH001, an agonistic monoclonal antibody against BTN3A1 developed in the course of the present study, disrupts the interaction between the BTN3A1 and BTN2A1 ectodomains. TH001 has a slightly higher binding affinity for BTN3A than the clinical candidate ICT01^23^ (Extended Data Fig. 4d), with superior performance over ICT01 for enhancing the responsiveness of Vγ9Vδ2 T cells for cancer cells (Extended Data Fig. 4e). Through cryo-EM analysis, we determined the structure of TH001-Fab in complex with HMBPP-primed BTN multimer at a resolution of 3.34 Å (Extended Data Fig. 1b, Extended Data Fig. 5 and Extended Data Table 2). TH001 engages both BTN3A1 and BTN3A2 within the heterodimer, burying approximately 800 Å² on each BTN3A molecule (Fig. 2b). TH001 occupies more epitopes on the BTN molecule than the antibody 20.1 (Fig. 2c, d). Notably, TH001 does not disrupt the association between the BTN3A and BTN2A1 ectodomains (Fig. 2b).

Finally, our two primed BTN multimer structures—neither of which is engaged with the Vγ9Vδ2 TCR—together reveal insights about the influence of T cell activators and suggest a plausible hypothesis about the structural changes likely to occur upon binding with the Vγ9Vδ2 TCR. That is, our structures support that the BTN3A–BTN2A1 ectodomain interaction persists even when these proteins are being primed for T cell activation by diverse molecules (*e.g.*, the natural pAg molecule HMBPP or the highly active BTN3A1 agonistic antibody TH001). And this persistence of the BTN3A–BTN2A1 ectodomain interaction, when viewed alongside previous structural evidence about the BTN2A1–Vγ9Vδ2 TCR binding interface, anticipates that the process of Vγ9Vδ2 TCR engagement must somehow cause dissociation of the BTN3A–BTN2A1 ectodomain interaction.

### Cryo-EM structure of the Vγ9Vδ2 TCR-engaged BTN3A1-BTN3A2–BTN2A1 complex

We next investigated how the Vγ9Vδ2 TCR engages the BTN3A1-BTN3A2–BTN2A1 complex to mediate T cell activation. We successfully assembled the Vγ9Vδ2 TCR (clone G115) in complex with HMBPP-primed BTN multimer. We conducted cryo-EM analysis and obtained the final reconstructions revealing the extracellular region at 3.21 Å resolution and the intracellular TM/JM/B30.2 regions at 4.51 Å resolution (Extended Data Fig. 2 and Extended Data Table 1).

In the TCR-disengaged state, the ectodomains of BTN3A2 and BTN2A1 remain associated. Upon TCR engagement, the width of the extracellular region remains almost unchanged at 140 Å (Fig. 1b and Fig. 3b), whereas the extracellular and JM regions of the BTN molecules undergo substantial rotational movements. This is most pronounced for BTN2A1, which undergoes a rigid-body shift, effectively “opening the pliers” collectively formed by BTN3A1-BTN3A2–BTN2A1 (Fig. 3c). In this “pliers-like arrangement”, BTN2A1 engages the lateral surface of the TCR, while BTN3A2 occupies its apical surface. Notably, this “pliers-like gripping” mechanism for dual ligand recognition revealed in our study contrasts sharply with previously reported αβ TCR–ligand interactions^24–26^ (Extended Data Fig. 6).

**Fig. 3.**
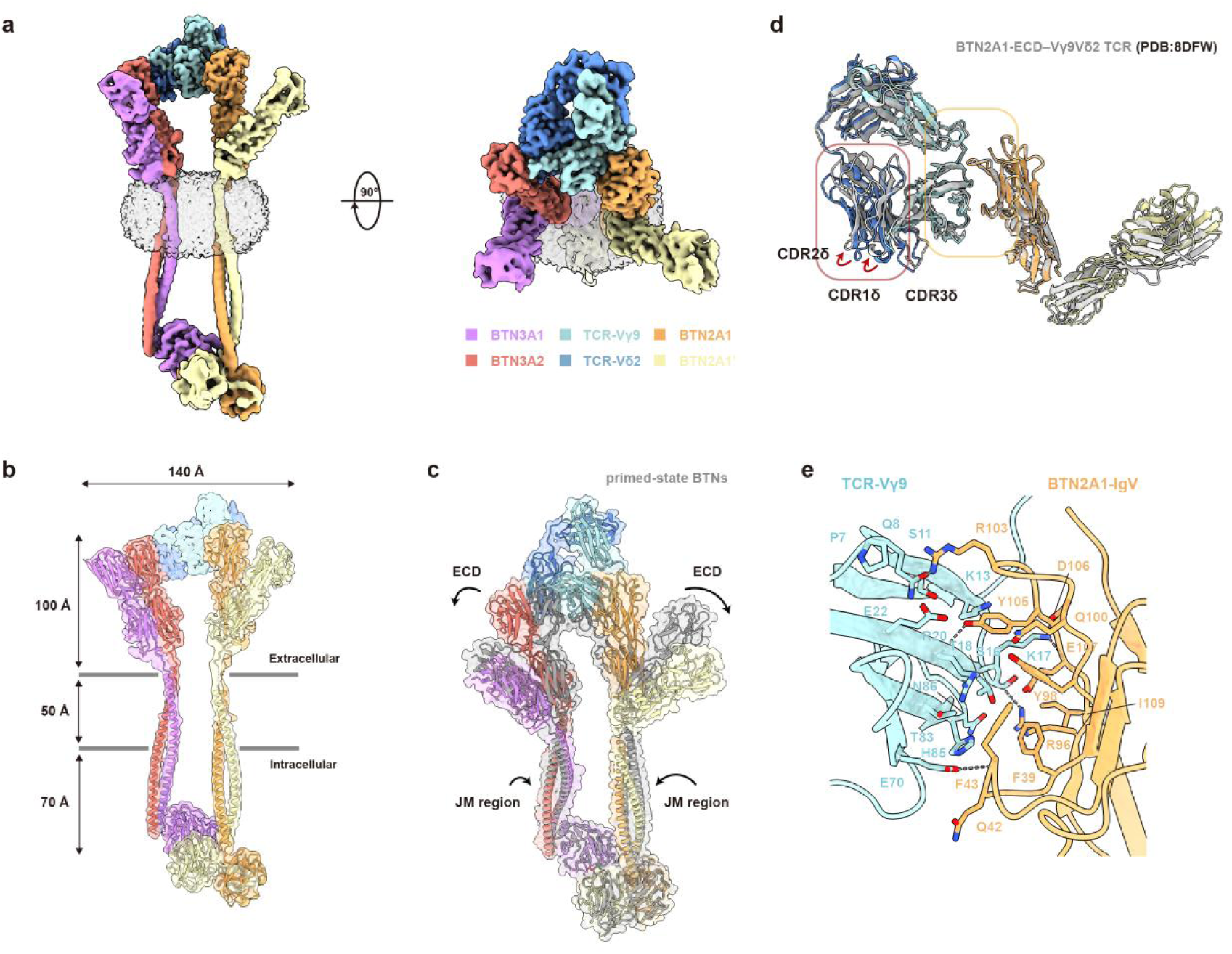
Cryo-EM structure of the Vγ9Vδ2 TCR-engaged BTN3A1-BTN3A2–BTN2A1 complex. **a,** Cryo-EM reconstruction of the complex. BTN3A1, BTN3A2, TCR**-**Vγ9, TCR**-**Vδ2, and the two chains of BTN2A1 are shown in purple, salmon, light blue, blue, yellow, and light yellow, respectively. The detergent micelles are depicted in transparent gray. Two perpendicular views are shown. **b,** The TM/JM regions of the BTN3A1**-**BTN3A2 heterodimer and the BTN2A1 homodimer form coiled-coil structures. Models of BTN molecules and the cryo-EM reconstruction are shown. **c,** The BTN3A1**-**BTN3A2**–**BTN2A1 complex (grey) undergoes conformational change upon Vγ9Vδ2 TCR engagement. Alignment was performed based on the B30.2 domains. **d,** Interaction mode between the Vγ9Vδ2 TCR and BTN2A1 resembles a previously determined crystal structure (gray, PDB ID: 8DFW). The yellow and red boxes indicate the BTN2A1 and BTN3A2 binding sites. **e,** Interactions between the Vγ9Vδ2 TCR and BTN2A1.

An intriguing finding from our TCR-engaged complex is that BTN3A2 interacts with the apical surface of the Vγ9Vδ2 TCR, primarily engaging with the Vδ2 chain and partially with the Vγ9 chain. This observation aligns with earlier crystal structure of the BTN2A1–Vγ9Vδ2 TCR complex (PDB 8DFW), where this surface was found to be unoccupied^18^. In parallel, the interaction between TCR and BTN2A1 mirrors previous findings^18^: the ABED face of Vγ9 interacts with the CFG face of BTN2A1, burying an area of approximately 500 Å² on BTN2A1 (Fig. 3d).

BTN2A1 primarily engages Vγ9Vδ2 TCR through its CC′-loop and G-strand, with five relevant hydrogen bonds: between Gln8γ and Arg103^BTN2A^^1^, Ser16γ and Arg96^BTN2A^^1^, Lys17γ and Glu107^BTN2A^^1^, Arg20γ and Tyr105^BTN2A^^1^, as well as Glu70γ and Phe43^BTN2A^^1^. Additional Vγ9 interactions of note including a salt bridge Lys17γ and Asp106^BTN2A^^1^ and a cation–π interaction between Arg20γ and Phe43 ^BTN2A^^1^ (Fig. 3e).

### Interactions between the Vγ9Vδ2 TCR and butyrophilins

Previous studies have established that BTN2A1 is a ligand for the Vγ9Vδ2 TCR^12,13^, but it has remained unclear whether BTN3A (or perhaps some other accessory protein) also serves as a ligand^18,27^. The involvement of BTN3A in this interaction has been particularly contentious, with several studies failing to detect a direct interaction^9,16,28^. Shedding light on the controversy about direct TCR engagement with BTN3A2, our high-resolution cryo-EM structure definitively establishes that BTN3A2 does function as *bona fide* a ligand for the Vγ9Vδ2 TCR (Fig. 3a, b). Extending beyond simple ligand binding, our structure also reveals a conformational change in the Vγ9Vδ2 TCR that occurs upon BTN3A2 binding (Fig. 3d and Fig. 4b). BTN3A2 induces a downward shift in the Vδ2 chain, which mediates the lion’s share of BTN3A2–TCR interactions. Specifically, complementarity-determining regions (CDRs) and hypervariable (HV) loops of the Vδ2 chain—specifically CDR1δ, CDR2δ, CDR3δ, and HV4δ—participate in the interactions with the BTN3A2 ligand that drive the TCR’s conformational change (Fig. 4a). The two chains of TCR bury approximately 700 Å² on BTN3A2.

**Fig. 4.**
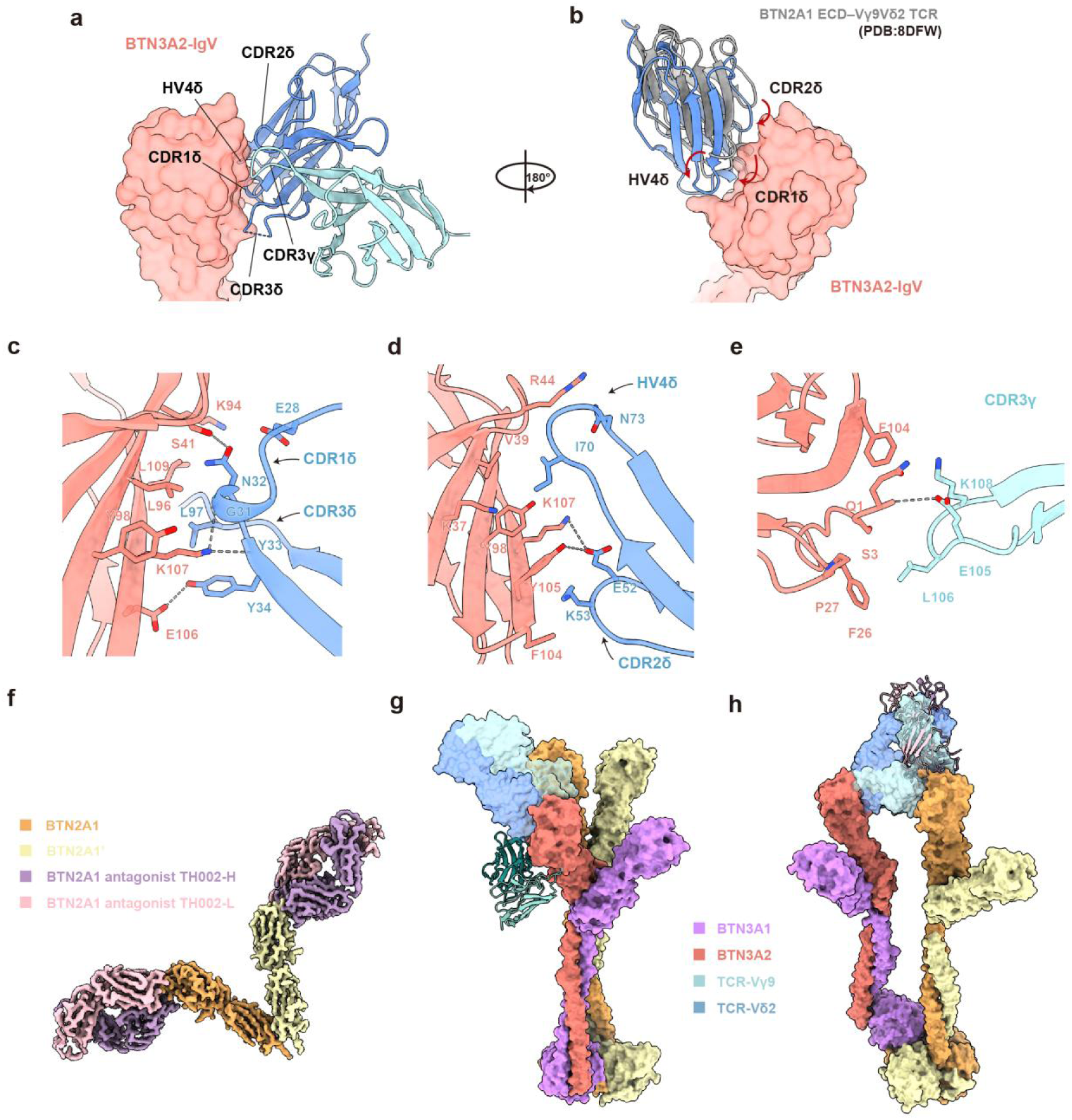
Interaction between the Vγ9Vδ2 TCR and butyrophilins. **a,** Overview of the interactions between the Vγ9Vδ2 TCR and BTN3A1-BTN3A2. The loops involved in the interaction are indicated by black lines. The IgV domain of BTN3A2 is shown as a surface representation in salmon. **b,** A zoomed-in and flipped view in the red box of Fig. 3d. The Vγ9Vδ2 TCR undergoes a conformational change upon binding to BTN3A1-BTN3A2. The IgV domain of BTN3A2 is shown as a surface representation in salmon. TCR**-**Vδ2 before and after BTN3A binding are shown in gray and blue, respectively. **c,** Interactions of CDR1δ and CDR3δ with BTN3A2. Dashed lines indicate polar interactions. **d,** Interactions of CDR2δ and HV4δ with BTN3A2. **e,** Vγ9 chain interaction with BTN3A2. **f,** Cryo-EM reconstruction of the BTN2A1–TH002-Fab complex. The two BTN2A1 molecules are shown in two shades of yellow. The heavy light chains of TH002 are shown in violet and pink. **g-h,** Superimposed model of the agonist antibody TH001 **(g)** or the antagonistic antibody TH002 **(h)** on the TCR-engaged BTN multimer. The complex is shown as a surface representation; TH001 and TH002 are shown as ribbons.

In CDR1δ, the main chain groups of Gly31δ and Tyr33δ form hydrogen bonds with Lys107^BTN3A^^2^, while the side chains of Asn32δ and Tyr34δ respectively establish hydrogen bonds with Ser41^BTN3A^^2^ and Glu106^BTN3A^^2^ (Fig. 4c). CDR2δ features a salt bridge between Glu52δ and Lys107^BTN3A^^2^, alongside hydrogen bonds with Tyr105^BTN3A^^2^, and Lys53δ engages in van der Waals interactions with Phe104^BTN3A^^2^ and Tyr105^BTN3A^^2^ (Fig. 4d). In CDR3δ, Leu97δ is sandwiched between Leu96^BTN3A^^2^ and Lys107^BTN3A^^2^ (Fig. 4c). And in HV4δ, Ile70δ participates in hydrophobic interactions and van der Waals interactions with Val39^BTN3A^^2^ and Tyr98^BTN3A^^2^ (Fig. 4d). Finally, note that the Vγ9 chain is also involved in the interaction with BTN3A2, primarily through its CDR3γ loop, for which Glu105γ forms a hydrogen bond with Gln1^BTN3A^^2^, while Lys108γ interacts with Phe104^BTN3A^^2^ (Fig. 4e).

Notably, there is obvious overlap between residues involved in the BTN3A2–BTN2A1 ectodomain interface and the BTN residues comprising the epitope for Vγ9Vδ2 TCR engagement (Extended Data Fig. 4a, b). By contrast, eight BTN3A2 residues are unique involved in TCR engagement (*i.e.*, are not involved in BTN3A2–BTN2A1 interface): Asn1, Phe2, Ser3, Phe26, Pro27, Lys37, Val39, and Phe104 (Extended Data Fig. 4a). The BTN2A1 residues unique to TCR engagement include Ile3, Arg96, Asp106, and Ile109 (Extended Data Fig. 4b).

Our clarification that the Vγ9Vδ2 TCR recognizes dual ligands, BTN3A2 and BTN2A1, opens the door for further exploration and structural characterization of functional butyrophilin antibodies. For instance, the BTN3A agonist antibody TH001 occupies the Arg44–Val46, Glu55, Glu57–Pro63, Arg65–Gly66, Ser69, and His85–Asn86 epitopes without interfering in BTN2A1–Vγ9Vδ2 TCR binding (Fig. 4g and Extended Data Fig. 8a). We also explored the mode of action for an antagonistic antibody against BTN2A1 that we developed in the present study using hybridoma technology (designated TH002). We assembled the BTN2A1–TH002-Fab complex and determined its cryo-EM structure at a resolution of 3.27 Å (Extended Data Fig. 1c, Extended Data Fig. 7 and Extended Data Table 2). In this structure, two TH002-Fab molecules symmetrically bind to both sides of the BTN2A1 V-dimer (Fig. 4f), burying an area of approximately 920 Å² on each BTN2A1 molecule. This interaction involves residues Gln1–Ile3, Leu25–Pro27, Lys29, Asp33–Met34, Lys51, Arg96, and Phe99–Ile109 (Extended Data Fig. 8b). The binding of TH002 effectively blocks the binding sites for the Vγ9Vδ2 TCR (Fig. 4h), thus explaining its antagonistic behavior (Extended Data Fig. 8c).

## DISCUSSION

The cryo-EM structures of the BTN3A1-BTN3A2–BTN2A1 complex, including those primed with HMBPP, an anti-BTN3A1 agonistic antibody (TH001), or engaged with the Vγ9Vδ2 TCR, reveal a distinctive mode of TCR activation. Upon TCR engagement, substantial conformational changes occur in the BTN multimers, leading to the dissociation of the BTN3A2 and BTN2A1 ectodomains and to formation of a “pliers-like grip” on the TCR.

This “pliers-like gripping” mechanism explains how different classes of Vγ9Vδ2 TCR activators, such as pAgs and BTN3A1 agonistic antibodies, operate. Consider that pAgs^15,29^ and BTN3A antibodies^30^ may promote clustering of BTN proteins, thereby enhancing their avidity for TCR engagement. The activators may facilitate “pliers-like gripping” in a thermodynamically favorable manner. HMBPP can induce an “inside-out” conformational change in BTN proteins, weakening their extracellular associations^15^, which could enhance the “pliers-like grip” on the TCR. Conversely, BTN3A1 agonistic antibodies stabilize the conformation of BTN3A, reducing entropy loss. This stabilization aligns with observations that BTN3A1 agonistic antibodies enhance the binding affinity of BTN3A1 for the Vγ9Vδ2 TCR^18^.

The structural similarity among BTN3A1, BTN3A2, and BTN3A3 have long suggested that BTN3A1 homodimers or BTN3A1–BTN3A3 heterodimers may engage the Vγ9Vδ2 TCR. However, previous efforts to identify BTN3A1 as a direct TCR ligand were unsuccessful^9,16,28^ (*i.e.*, no interaction was detected). Our cryo-EM analysis brings clarity by showing that a conformational change in the TCR makes it accessible for BTN3A binding. Uldrich *et al.* (2024) reported that lysine 53 (K53) in the TCR negatively regulates Vγ9Vδ2 T cell activation, and a single mutation of K53 to alanine (A) significantly enhances the TCR’s capacity to engage BTN3A, leading to spontaneous T cell activation^18^. Notably, K53 is located within the CDR2δ region, which we found to participate in the interactions with the BTN3A ligand that drive the TCR’s conformational change through the piler-like gripping mechanism. It is likely that the K53A mutation promotes a similar conformational shift, making the TCR more conducive to BTN3A1 binding. This may involve a structural rearrangement that exposes previously hidden regions of the TCR or that modify the interaction interface.

Previous studies have implicated various pairs of BTN and BTNL/Btnl members of the B7 superfamily in regulating γδ T cell activation in humans and mice^9^. For instance, BTNL3–BTNL8 heterodimers are responsible for activating Vγ4 γδ T cells in the human gut^31,32^. In mice, Btnl1–Btnl6 heterodimers enhance the activation of gut-resident Vγ7 γδ T cells, and Skint1–Skint2 heterodimers enhance activation of skin-resident Vγ5Vδ1 dendritic epidermal T cells^31,33–35^. Our previous study using both structural data and activation assays with mutant BTN variants demonstrated that Vγ9Vδ2 T cells use both BTN3A1 and BTN2A1 for efficiently detecting intracellular metabolites^15^. The present study establishes that BTN3A2 and BTN2A1 are both ligands for the Vγ9Vδ2 TCR. Our elucidation of the “pliers-like gripping” mechanism raises the possibility that such dual-ligand recognition may not be unique to the Vγ9Vδ2 TCR. It is plausible that other γδ TCRs may employ a similar mechanism for antigen recognition and γδ T cell activation.

## Materials and Methods

### Cell lines

#### HEK293F cells

HEK293F cells (Thermo) were cultured in FreeStyle 293 Expression Medium (Thermo) at 37°C with 5% CO₂ and 55% humidity using a humidified shaker for protein purification.

#### MIA PaCa-2 cells

MIA PaCa-2-luciferase cells were generated by infecting MIA PaCa-2 cells with lentivirus carrying the luciferase transgene and BTN2A^−/−^ MIA PaCa-2 (BTN2A1/BTN2A2 KO) cells were generated by disrupting the *BTN2A1* and *BTN2A2* genes in MIA PaCa-2 cells using CRISPR^15^. MIA PaCa-2 cells were cultured in Dulbecco’s Modified Eagle’s Medium (DMEM) (Gibco) supplemented with 10% heat-inactivated FBS and 1% penicillin–streptomycin (Beyotime) at 37°C in a 5% CO₂ atmosphere.

#### CHO-K1 cells

CHO-K1 cells, a subclone derived from the parental CHO cell line obtained from an ovarian biopsy of an adult Chinese hamster, were purchased from ATCC and cultured in Ham’s F12K (Kaighn’s) medium (HyClone) supplemented with 10% heat-inactivated FBS and 1% penicillin-streptomycin at 37°C in 5% CO_2_. To generate cells overexpressing BTN3A1 or BTN2A1, CHO-K1 cells were infected with lentivirus carrying the *BTN3A1* or *BTN2A1* transgenes; successfully transduced cells selected using 1 mg/mL puromycin.

#### Human Vγ9Vδ2 T cells

All procedures involving Vγ9Vδ2 T cells complied with the Institutional Review Board guidelines of Tsinghua University, with written informed consent obtained from all participants, as per the Declaration of Helsinki. The protocol was approved by the Institutional Review Board (project nos. 20170004 and 20170007). Peripheral blood mononuclear cells (PBMCs) were isolated from healthy donors via density-gradient centrifugation using Ficoll-Hypaque (GE Healthcare). Cells were cultured at 2 × 10⁶ cells/ml in RPMI 1640 (Gibco) supplemented with 10% FBS, 1% penicillin-streptomycin, 150 U/ml human recombinant IL-2 (PeproTech), 1% MEM non-essential amino acids (Gibco), 2 mM L-glutamine (Beyotime), 50 mM β-mercaptoethanol (Amresco), and 5 µM zoledronate (Energy Chemical) at 37°C in a 5% CO₂ atmosphere. The medium was replaced with fresh IL-2 every 3 days. Vγ9Vδ2 T cells were harvested on day 11, achieving >90% purity, and were stored for future use.

### Protein purification

#### Full-length butyrophilin proteins

Genes encoding BTN2A1 (UniProtKB: Q7KYR7) and BTN3A1 (UniProtKB: O00481) were cloned into the pcDNA3.1 vector with a C-terminal Twin-strep tag. BTN3A2 (UniProtKB: P78410) was cloned into the pcDNA3.1 vector with a C-terminal Flag tag. HEK293F cells were grown to a density of 2×10^6^ cells per milliliter and transfected with plasmids encoding BTN3A1 and BTN3A2 at 1:1 ratio or BTN2A1 alone using 40 kDa linear polyethylenimine (PEI; Polysciences). For one liter of cell culture, 2 mg of plasmids was mixed with 4 mg of PEI in 50 ml fresh medium, and incubated for 30 min before transfection. Transfected cells were cultured for another 60 hours, then collected by centrifugation.

For full-length BTN2A1 or BTN3A1-BTN3A2 protein purification, cell pellets were resuspended, and membranes were solubilized in a buffer containing 25 mM HEPES pH 7.4, 150 mM NaCl, 1% (w/v) lauryl maltose neopentyl glycol (LMNG; Anatrace) and 0.1% (w/v) cholesteryl hemisuccinate Tris salt (CHS; Anatrace) supplemented with 1 mM phenylmethylsulfonyl fluoride (PMSF) and 1 × protease inhibitor cocktail (Bimake). The suspension was rotated for 3 hours at 4℃ followed by centrifugation at 8,000 rpm for 40 minutes and ultracentrifugation at 40,000 rpm for 1 hour. After removing debris, the resulting supernatant was incubated with Strep-tactin resin (GenScript) for 2 hours. The resin was washed with buffer containing 25 mM HEPES pH 7.4, 150 mM NaCl, 0.5% (w/v) LMNG, and 0.05% (w/v) CHS, followed by washing with size-exclusion chromatography (SEC) buffer containing 25 mM HEPES pH 7.4, 150 mM NaCl, 0.001% (w/v) LMNG, 0.0001% (w/v) CHS, and 0.00033% glyco-diosgenin (GDN; Anatrace). The BTN2A1 or BTN3A1-BTN3A2 complex was then eluted with the SEC buffer supplemented with 10 mM desthiobiotin (IBA Lifesciences), and further purified by SEC (Superose 6 Increase 10/300 GL, Cytiva) in SEC buffer. The protein fractions were monitored at all stages of purification using SDS-PAGE and visualized by Coomassie blue staining. The resulting proteins (after concentration) were flash frozen and stored at −80℃.

#### Ectodomains of Vγ9Vδ2 TCR, BTN2A1 and BTN3A1

For expression of Vγ9Vδ2 TCR (G115) ectodomain, genes encoding the T-cell receptor γ9 chain (GenBank: CAA51165.1, residues 21-262) and δ2 chain (PIR: S33439, residues 20-248) were cloned into the pcDNA3.1 vector with a N-terminal human IL-10 signal peptide (UniProtKB: P22301, residues 1-18) and a C-terminal acidic or basic leucine zipper, followed by an 8×His tag in the case of γ9 chain. For expression of BTN2A1 and BTN3A1 ectodomains, genes encoding the BTN2A1 or BTN3A1 ectodomain was cloned into the pcDNA3.1 vector with a C-terminal 8×His tag. HEK293F cells were grown to a density of 1.0 × 10^6^ cells per milliliter and transfected with the plasmids encoding γ9 and δ2 at 1:2 ratio or BTNs’ ectodomain alone using PEI. For one liter of cell culture, 1 mg of plasmids was mixed with 2 mg of PEI in 50 ml fresh medium, and incubated for 30 min before transfection.

For the above proteins, the medium supernatant was collected by centrifugation after 4 days of transfection. Then the supernatant was concentrated and exchanged into the binding buffer (25 mM HEPES pH 7.4, 150 mM NaCl) with a Hydrosart Ultrafilter (Sartorius). The proteins were isolated by Ni-NTA affinity chromatography, eluted with the binding buffer supplemented with 500 mM imidazole (Sigma-Aldrich), and further purified by SEC (Superose 6 Increase 10/300 GL, Cytiva) in the same SEC buffer as that for full-length butyrophilin proteins. The purified proteins after concentration were flash frozen and stored at −80℃.

#### Production of monoclonal antibodies

Mouse antibodies against human BTN3A1 (TH001) and BTN2A1 (TH002) were generated by immunizing mice with recombinant human BTN3A1-Fc or BTN2A1-Fc fusion proteins. A total of 50 μg of each protein, mixed with Freund’s adjuvant (Sigma), was intramuscularly injected into mice five times at three-week intervals. The antibody titer in mouse serum was assessed seven days after the final immunization. Mice with the highest specific antibody titers for BTN3A1 or BTN2A1 were euthanized, and their spleens were collected. A single-cell suspension of splenocytes was prepared using a 70 μm filter, followed by PEG-induced fusion with myeloma cells to generate hybridomas, which were cloned in HAT as previously described^36^. All hybrid clone supernatants were screened for antibody detection using ELISA with specific recombinant protein coating and flow cytometry with CHO-K1 cells or CHO-K1 cells overexpressing BTN3A1 (CHO-3A1) or BTN2A1 (CHO-2A1). The hybridoma cell lines secreting monoclonal antibodies (mAbs) against BTN3A1 or BTN2A1 were obtained through three rounds of subcloning and were subsequently injected into sensitized mice to produce ascitic fluid^36^. The antibodies were purified from the ascitic fluid, and their purity was analyzed by SDS-PAGE. The antigen-binding fragments (Fab) of TH001 and TH002 were produced through papain cleavage, as described previously^37^.

#### Cryo-EM sample preparation and data collection

For the human BTN3A1**-**BTN3A2**–**BTN2A1 complex, purified full-length BTN2A1, full-length BTN3A1**-**BTN3A2 and Vγ9Vδ2 TCR were mixed at a 1:1:1 molar ratio and incubated on ice for 2 h. HMBPP was then added at a final concentration of 1 μM. The BTN3A1**-**BTN3A2–BTN2A1–TH001-Fab complex was obtained similarly. For the BTN2A1–TH002**-**Fab complex, purified full-length BTN2A1 and TH002-Fab were mixed at a 1:1 molar ratio and incubated on ice for 2 h without HMBPP. The protein mixtures were loaded onto a Superose 6 Increase column and eluted with the aforementioned SEC buffer. The peak fractions corresponding to the complex were collected and concentrated to 4 mg/mL for cryo-EM sample preparation.

Cryo-grids were prepared using the Vitrobot Mark IV (FEI). Briefly, a 4 μL aliquot of protein complex was applied to glow-discharged holy-carbon gold grids (Quantifoil, R1.2/1.3, 300-mesh) or graphene oxide (GO)-coated holey-carbon gold grids (EMR, R1.2/1.3, 400-mesh). Grids were blotted for 3.0 s and quickly plunged into liquid ethane under 6℃ and 100% humidity.

Data collection was carried out on a 300 kV Titan Krios G4 electron microscope equipped with a Falcon 4 camera. Movie stacks were recorded using the EPU software (Thermo Scientific) at a magnification of 130,000×, corresponding to a calibrated pixel size of 0.95 Å at object scale, and with a preset defocus range from −1.3 to −1.7 μm. Statistics for data collection are summarized in Extended Data Table 1 and 2.

#### Cryo-EM data processing

The flowcharts for data processing of the BTN3A1-BTN3A2–BTN2A1, BTN3A1-BTN3A2–BTN2A1–TH001-Fab and BTN2A1–TH002-Fab complexes are respectively presented in Extended Data Fig. 2, 5 and 7. The datasets were processed using cryoSPARC^38^.

For the primed BTN3A1-BTN3A2–BTN2A1 and TCR-engaged and structures, a total of 17,303 movie stacks were collected. Raw movies were motion-corrected using Patch motion correction, and dose-weighted micrographs were subjected to Patch CTF estimation. Micrographs with ice contamination or exhibiting an estimated CTF resolution of worse than 4 Å were discarded. A subset of 2,000 micrographs was subjected to blob picking and multiple rounds of 2D classification to generate good 2D class averages. Particle picking was then performed by template picking and topaz picking to generate a dataset of 11,873,703 particles. Extracted particles were binned by a factor of 2 and subjected to multiple rounds of 2D classification to remove junk particles. Selected particles were re-centered and re-extracted without binning. Then, a subset of 500,000 particles was subjected to Ab-Initio reconstruction and heterogeneous refinement to further remove junk particles. Particles of good classes were subjected to non-uniform (NU) refinement and 3D classification to identify potential multiple states. 3D references of the TCR-engaged state and the primed state were further generated using NU refinement and local refinement.

Preliminarily screened particles from the whole dataset were subjected to multiple rounds of heterogeneous refinement with three bad references generated from junk particles and two good references of TCR-engaged state and primed state. Subsequently, particles of the TCR-engaged state and the primed state with duplicates removed were subjected to NU refinement and local refinement, yielding a global 3.68 Å map of the TCR-engaged state and a global 4.13 Å map of the primed state.

To retrieve good particles in bad classes, the method of seed-facilitated 3D classification^39^ was performed by using the previous reconstruction particles of both states as good seeds. Raw particles were divided into subsets and subjected to multiple rounds of heterogeneous refinement with good and bad references, until more than 97% of the remaining particles were classified into good classes. Good particles of each state were re-centered and re-extracted, then subjected to one round of heterogeneous refinement with resolution-gradient references. After removing duplicates, the resulting particles were subjected to NU refinement followed by global CTF refinement, yielding a global 3.22 Å map of the TCR-engaged state and a global 3.74 Å map of the primed state. To improve the local resolution of both states, local refinement was performed with a soft mask applied to the extracellular region or TM/JM/B30.2 regions of the complex. For the TCR-engaged state, the ultimate 282,479 particles resulted in a map of the extracellular region at 3.21 Å and a map of TM/JM/B30.2 regions at 4.51 Å. For the primed state, the ultimate 264,215 particles resulted in a map of extracellular region at 4.17 Å and a map of TM/JM/B30.2 regions at 3.58 Å.

For the BTN3A1-BTN3A2–BTN2A1–TH001-Fab structure, a total of 6,978 movies for samples on holey-carbon gold grids and 4,457 movies for samples on GO-coated holey-carbon gold grids were collected. In total, 3,940,120 and 3,697,802 raw particles were automatically extracted from the two datasets. Extracted particles were binned by a factor of 2 and subjected to multiple rounds of 2D classification to remove junk. Selected particles were re-centered and re-extracted without binning, and subjected to Ab-Initio reconstruction and heterogeneous refinement. A good initial model was obtained and the corresponding 479,276 particles were subjected to NU refinement, yielding a map with a resolution of 3.38 Å. Subsequently, local refinement was performed with a soft mask applied to the extracellular region of BTN molecules and the variable fragment (Fv) region of TH001-Fab, yielding a final map at 3.34 Å.

For the BTN2A1–TH002-Fab structure, the data processing was essentially consistent with the above. A total of 5,926,117 raw particles were automatically extracted from 6,679 micrographs. After multiple rounds of 2D classification, selected particles were re-extracted and subjected to Ab-Initio reconstruction followed by multiple rounds of heterogeneous refinement to remove junk. The resulting particles without duplicates were subjected to NU refinement, yielding a map at 3.50 Å. After several rounds of seed-facilitated 3D classification, the ultimate 271,629 particles were subjected to NU refinement, followed by local refinement with a soft mask applied to the extracellular region of BTN2A1 and TH002-Fab, yielding a final map at 3.26 Å.

The masks were created using UCSF ChimeraX^40^. All the resolution estimations were based on a Fourier shell correlation (FSC) of 0.143 criterion. Local resolution estimation jobs were performed in cryoSPARC.

#### Model building and refinement

Final maps were post-processed using EMReady^41^ to reduce noise and enhance protein signal. B-factors sharpened maps from cryoSPARC and EMReady maps were used to assist model building.

For the BTN3A1-BTN3A2–BTN2A1 complex, the post-processed map of the extracellular region and the map of TM/JM/B30.2 regions for the TCR-engaged state or the primed state were combined using UCSF ChimeraX. Local resolution estimation of the composite map was performed using the composite half maps. For model building of the TCR-engaged state, the models of BTN2A1 (PDB ID: 8DFW), BTN3A1 (PDB ID: 4F80), BTN3A2 (PDB ID: 4F8Q), and the Vγ9Vδ2 TCR (PDB ID: 8DFW) ectodomains from crystal structures, together with the BTN3A1-B30.2–HMBPP–BTN2A1-B30.2 model (PDB ID: 8JYE) and the TM/JM region models from Alphafold^42^, were docked into the composite map of the TCR-engaged state using UCSF Chimera. The primed state model was built similarly. For the BTN3A1-BTN3A2–BTN2A1–TH001-Fab and BTN2A1–TH002-Fab complexes, the model of TH001-Fab or TH002-Fab was predicted by Alphafold3 and docked into the corresponding density map. Models were manually adjusted using Coot^43^ and automatically refined using the real-space refinement in Phenix^44^. Figures were prepared with UCSF ChimeraX.

### Functional assays

#### Surface plasmon resonance

SPR experiments were conducted at 25°C on a Biocore 8K plus instrument (Cytiva) using PBS buffer containing 0.05% Tween 20. For the affinity assay of the butyrophilins with antibodies, the BTN3A1 and BTN2A1 ectodomains were respectively immobilized to 2381 and 2405 resonance units (RU) on a Biocore CM5 chip in 10 mM sodium acetate buffer (pH 5.5), at a concentration of 20 μg/mL. TH001 or TH002 were used as the analyte (diluted in running buffer), with concentrations ranging from 0.2 nM to 200 nM. After subtraction of data from the control flow cell and blank injections, interactions were analyzed with Biocore 8K insight evaluation software (Cytiva), and equilibrium dissociation constants (*K*_D_’s) were derived at equilibrium.

#### IFN-γ secretion assays of Vγ9Vδ2 T cells co-cultured with BTN2A^−/−^ MIA PaCa-2 cells

BTN2A^−/−^ MIA PaCa-2 (BTN2A1/BTN2A2 KO) cells were plated in 96-well plates at a ratio of 1.0 × 10^4^ cells per well 1 day prior to transfection. All plasmids were transfected into MIA PaCa-2 cells using Lipofectamine 2000 reagent and then treated with zoledronate (10 μM) for 24 h. The medium was then aspirated and the cells were washed 4 times with PBS at room temperature. Vγ9Vδ2 T cells (1.0 × 10^4^ cells per well) were added to MIA PaCa-2 cells for co-culture, and the culture supernatant was collected after 16 h with an IFN human Uncoated ELISA Assay Kit (Invitrogen).

#### Functional assays of antibodies against BTN3A1 and BTN2A1

The function of antibodies against BTN3A1 and BTN2A1 was evaluated using Vγ9Vδ2 T cells co-cultured with luciferase-expressing MIA PaCa-2 cells. MIA PaCa-2 cells were seeded at a density of 1.0 × 10^4^ cells per well in 96-well plates overnight. For the anti-BTN3A1 mAb, the antibody or isotype control was diluted from the final maximum concentration of 10 μg/mL with 10-fold gradient dilution steps (for a total of 8 points) and incubated with the MIA PaCa-2 cells for 12 h. Following two washes with PBS to remove the antibody, purified Vγ9Vδ2 T cells were added at a 1:2 ratio and co-cultured for 24 h. For the anti-BTN2A1 mAb, the antibody or isotype control was diluted from the final maximum concentration of 50 μg/mL with 10-fold gradient dilution steps (for a total of 8 points) and incubated with the MIA PaCa-2 cells for 2 h. Subsequently, 2.0 × 10^4^ Vγ9Vδ2 T cells were added for additional 2 h and cells were then stimulated for 18 h with ICT01 mAb (1 μg/mL). Cell killing was measured using Stable-lite luciferase (Vazyme) according to the manufacturer’s instructions.

## Acknowledgement

This work was supported by the National Natural Science Foundation of China (grant numbers 82350109, 82341040, 81991492, 32325018, 82341210), the National Key Research and Development Program of China (grant numbers 2021YFC2302604, 2023YFC2306202), and the Beijing Natural Science Foundation (grant number 5244037). Additional support was provided by the Qidong-SLS Innovation Fund to J.X., Changping Laboratory, the Tsinghua-Peking University Center for Life Sciences, the Beijing Frontier Research Center for Biological Structure, and the Postdoctoral Fellowship Program of CPSF (grant numbers GZB20230364, 2024M751731). We extend our gratitude to the Protein Preparation and Identification Facilities at Tsinghua University for help with SPR data analysis and to the cryo-EM and high-performance computing platforms at Peking University and Changping Laboratory for assistance with data collection and computation.

## Author contributions

Y.Z. and J.X. conceptualized and designed the project. M.Z., N.C., Y.W., Y.Q., X.M., J. X. and X.C. conducted the experiments. Y.Q., M.Z., Y.W. and J.X. were responsible for protein purification. M.Z. and Y.W. performed the cryo-EM analysis. N.C. and X.Z. developed and evaluated the antibodies. N.C., Y.Q., and X.C. carried out the cell assays. X.M. performed SPR analysis. Y.Z. and J.X. supervised the research. Y.Z., J.X., N.C., X.M. and M.Z. contributed to writing the manuscript.

## Additional information

Correspondence and requests for material should be addressed to Yonghui Zhang or Junyu Xiao.

## Data and materials availability

Cryo-EM density maps of the HMBPP-primed BTN3A1-BTN3A2–BTN2A1 complex have been deposited in the Electron Microscopy Data Bank with accession codes EMD-61736 (composite map), EMD-61733 (focused map of extracellular region), EMD-61734 (focused map of TM/JM/B30.2 regions) and EMD-61735 (raw consensus map). Cryo-EM density maps of the Vγ9Vδ2 TCR-engaged BTN3A1-BTN3A2–BTN2A1 complex have been deposited in the Electron Microscopy Data Bank with accession codes EMD-61740 (composite map), EMD-61737 (focused map of extracellular region), EMD-61738 (focused map of TM/JM/B30.2 regions) and EMD-61739 (raw consensus map). Structural coordinates have been deposited in the Protein Data Bank with the accession codes 9JQQ (HMBPP-primed state) and 9JQR (TCR-engaged state).

Meanwhile, cryo-EM density maps of the BTN3A1-BTN3A2–BTN2A1–TH001-Fab and BTN2A1–TH002-Fab complexes have been deposited in the Electron Microscopy Data Bank with accession codes EMD-61732 and EMD-61724. Corresponding coordinates have been deposited in the Protein Data Bank with the accession codes 9JQP and 9JQ6. The other structural coordinates used in this study are available from PDB (8JYE, 8DFW, 4F80, 4F8Q, 7PHR, 2PO6 and 6PUC). Source data are provided with this paper.

## EXTENDED DATAFIGURES

**Extended Data Fig. 1.**
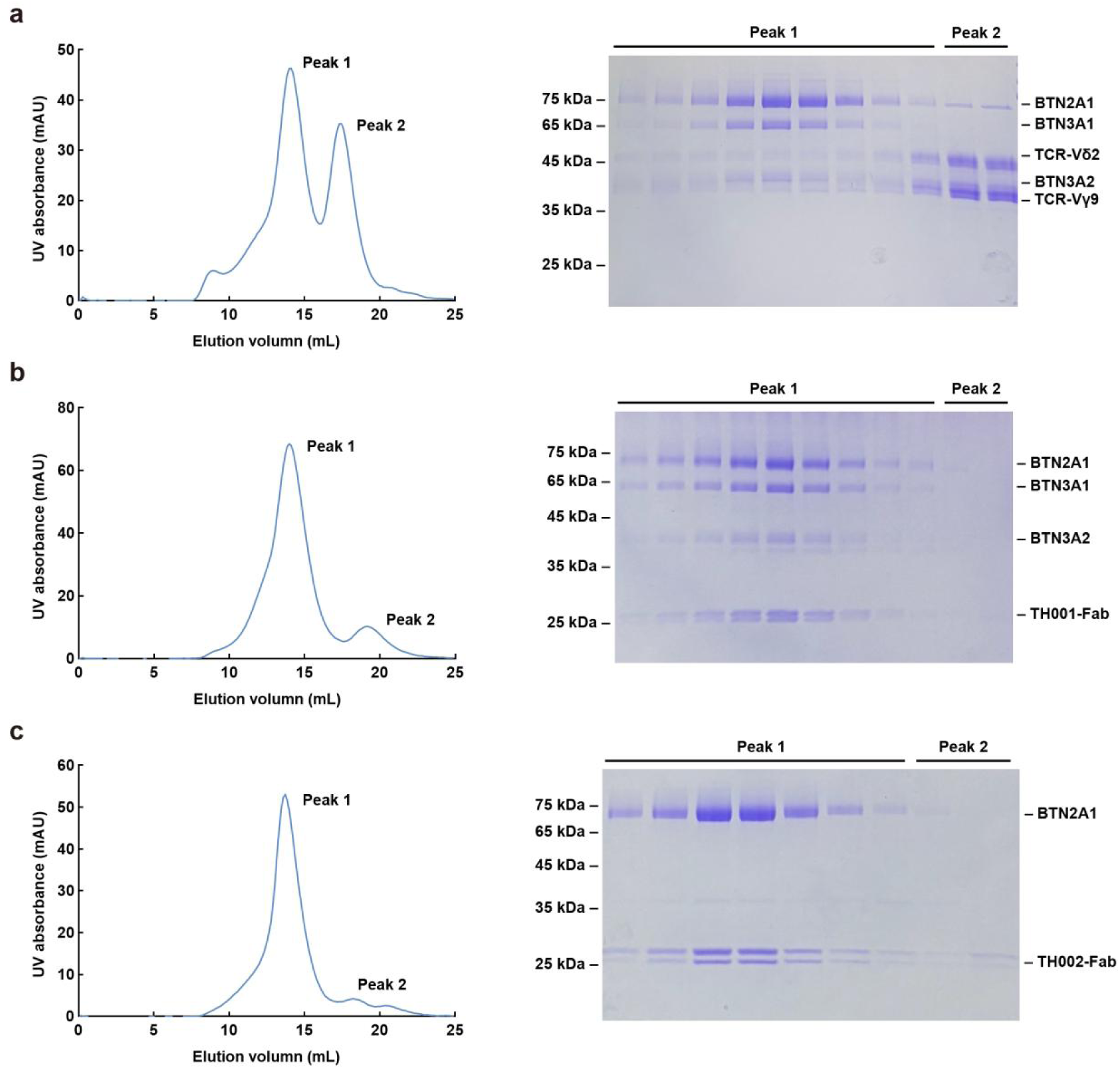
Purification of protein complexes for cryo-EM. **a.** Size-exclusion chromatography of the BTN3A1-BTN3A2–BTN2A1 complex on a Superose 6 Increase column (left) and corresponding SDS-PAGE analysis, visualized through Coomassie blue staining (right). All the purification experiments were repeated at least twice, with similar results. **b.** Size-exclusion chromatography of the BTN3A1-BTN3A2–BTN2A1–TH001-Fab complex on a Superose 6 Increase column (left) and corresponding SDS-PAGE analysis (right). **c.** Size-exclusion chromatography of the BTN2A1–TH002-Fab complex on a Superose 6 Increase column (left) and corresponding SDS-PAGE analysis (right).

**Extended Data Fig. 2.**
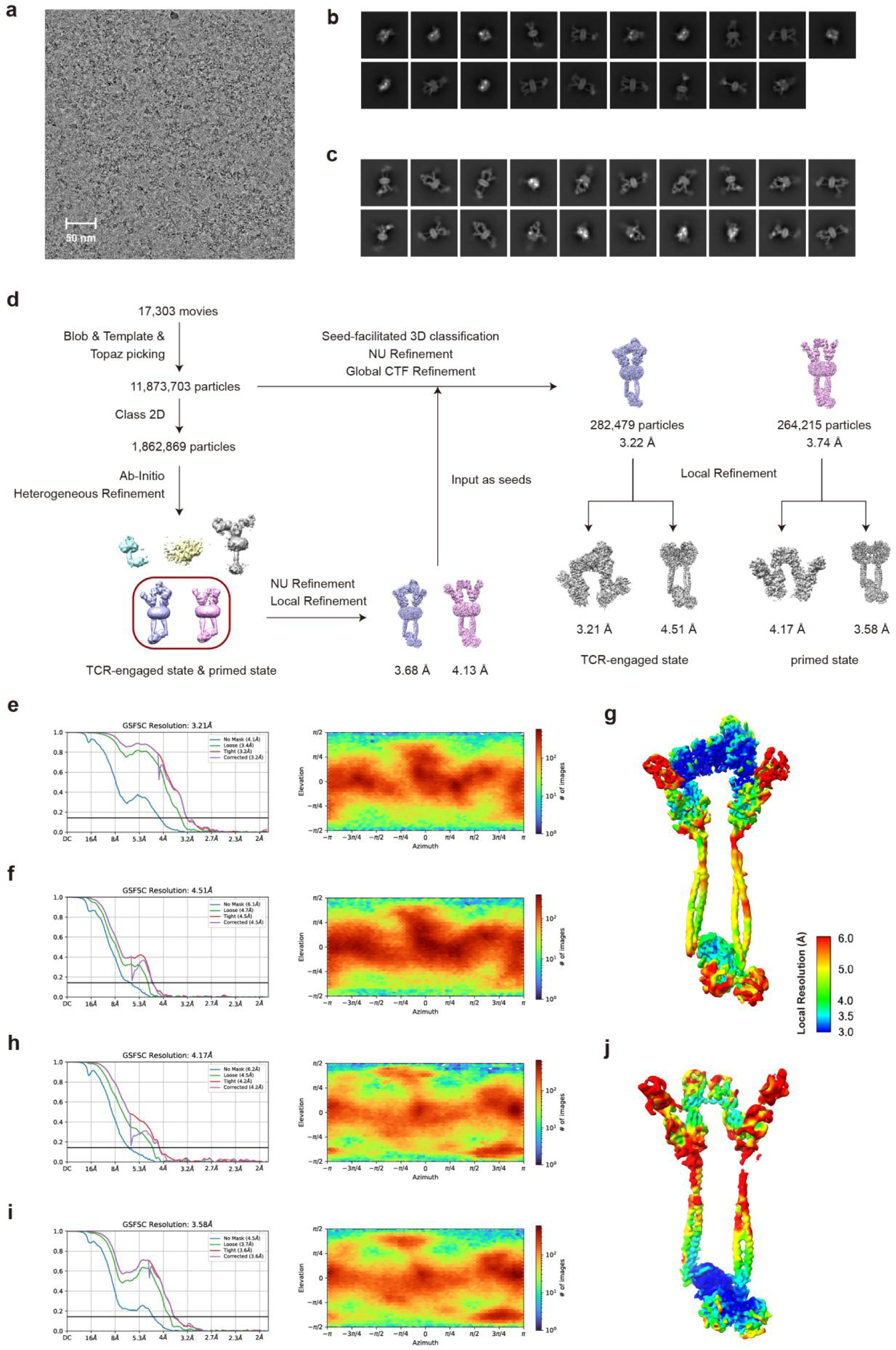
Cryo-EM analysis of the BTN3A1-BTN3A2–BTN2A1 complex. **a-c.** Representative cryo-EM micrograph (**a**) and 2D class averages of HMBPP-primed (**b**) or Vγ9Vδ2 TCR-engaged (**c**) BTN3A1-BTN3A2–BTN2A1 complex. Scale bar: 50 nm. **d.** Flowchart of the cryo-EM analysis for the BTN3A1-BTN3A2–BTN2A1 complex. **e-f.** Cryo-EM analysis for the reconstruction of the extracellular region and TM/JM/B30.2 regions in TCR-engaged state. Left: Gold standard Fourier shell correlation (GSFSC) curves of the density map. Right: Angular particle distribution heat map in the final round of local refinement. **g.** Local resolution estimation of the composite density map in TCR-engaged state. **h-i.** Cryo-EM analysis for the reconstruction of extracellular region and TM/JM/B30.2 regions in primed state. The order of the plots is the same as that in **e-f**. **j.** Local resolution estimation of the composite density map in the primed state.

**Extended Data Fig. 3.**
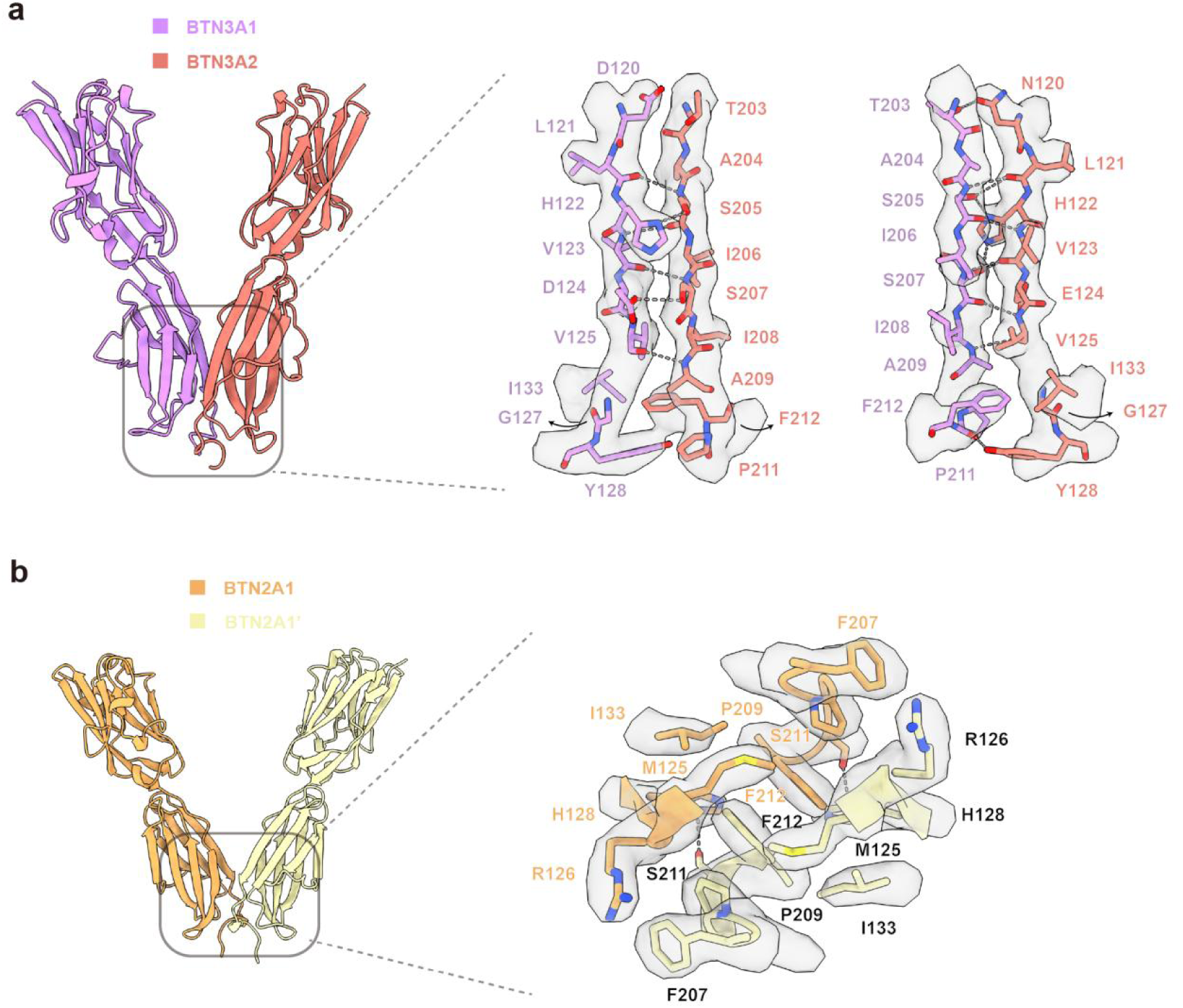
Interactions within the BTN3A1-BTN3A2 heterodimer and within the BTN2A1 homodimer. Interaction between the IgC domains of BTN3A1 and BTN3A2 **(a)** or between the IgC domains of two BTN2A1 molecules **(b)**. Models of BTN molecules (left) and residues within the interaction (right) are shown. The cryo-EM reconstructions are shown in transparent grey. Dashed lines indicate polar interactions.

**Extended Data Fig. 4.**
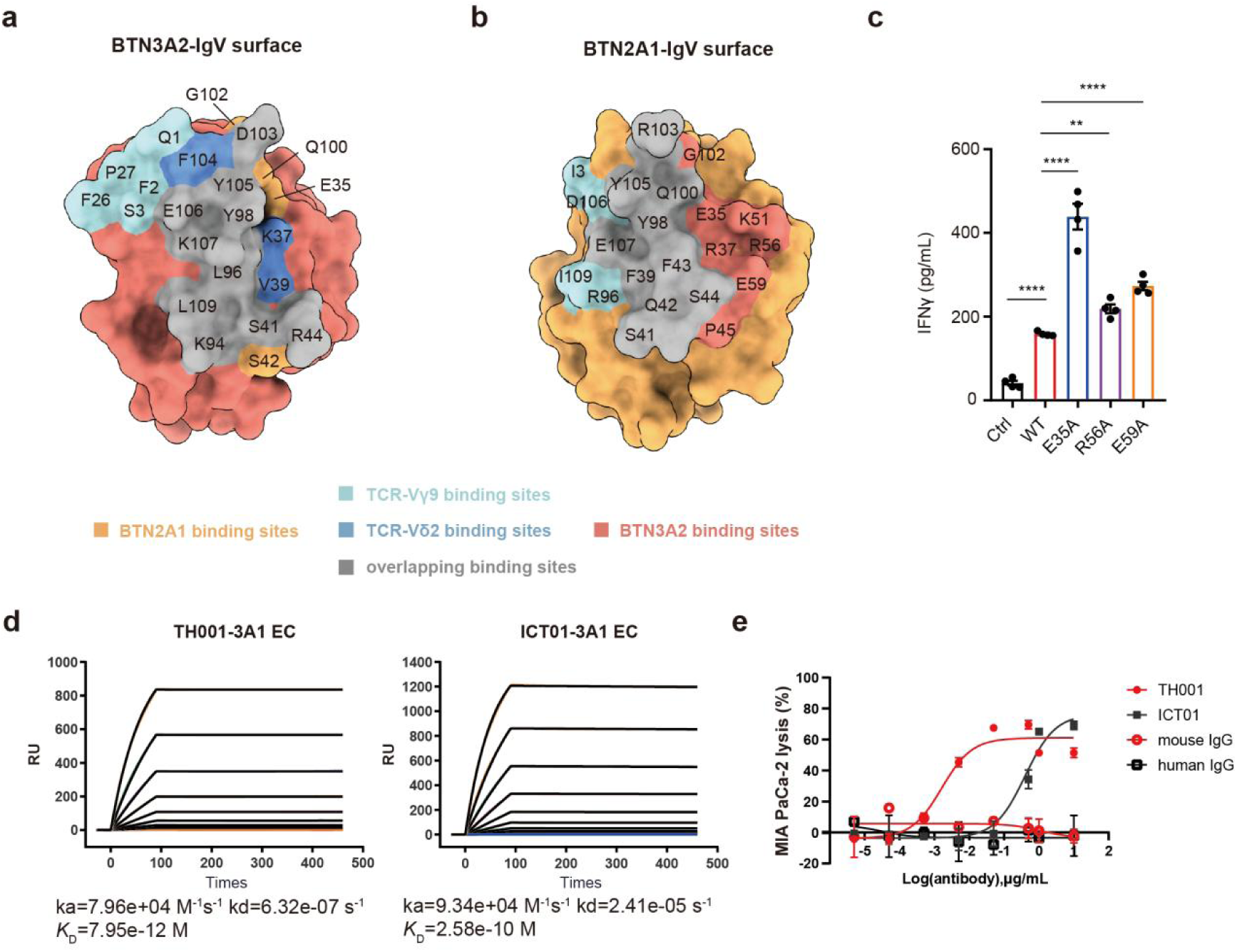
BTN ectodomain interactions. **a.** Surface representation of BTN3A2, depicting the regions that contact BTN3A2 and the Vγ9Vδ2 TCR. Unique binding sites of BTN2A1 and the Vγ9Vδ2 TCR are shown with the colors as indicted. Overlapping binding sites are shown in grey. **b.** Surface representation of BTN2A1 depicting the regions that contact BTN2A1 and the Vγ9Vδ2 TCR. **c,** Mutations in BTN2A1 IgV residues that prevent the BTN3A1-BTN2A1 ectodomain interaction enhance the responsiveness of Mia PaCa-2 cells (pretreated with zoledronate) to Vγ9Vδ2 T cells. *n*=4. Representative of three independent experiments. Data were analyzed with two-tailed unpaired t tests. NS, not significant, \*\**P* < 0.01, \*\*\**P* < 0.001 and \*\*\*\**P* < 0.0001. All error bars denote the SEM. **d,** SPR analysis indicating that the monoclonal antibody TH001 developed in the present study has a binding affinity for BTN3A comparable to the clinical candidate ICT01. **e,** Cytotoxicity of Vγ9Vδ2 T cells towards MIA PaCa-2 cells, pretreated with TH001 or ICT01 overnight. Mouse IgG (IgG1) or human IgG (IgG1) was used as the isotype control for TH001 or ICT01. Representative of three independent experiments. All error bars denote the SEM.

**Extended Data Fig. 5.**
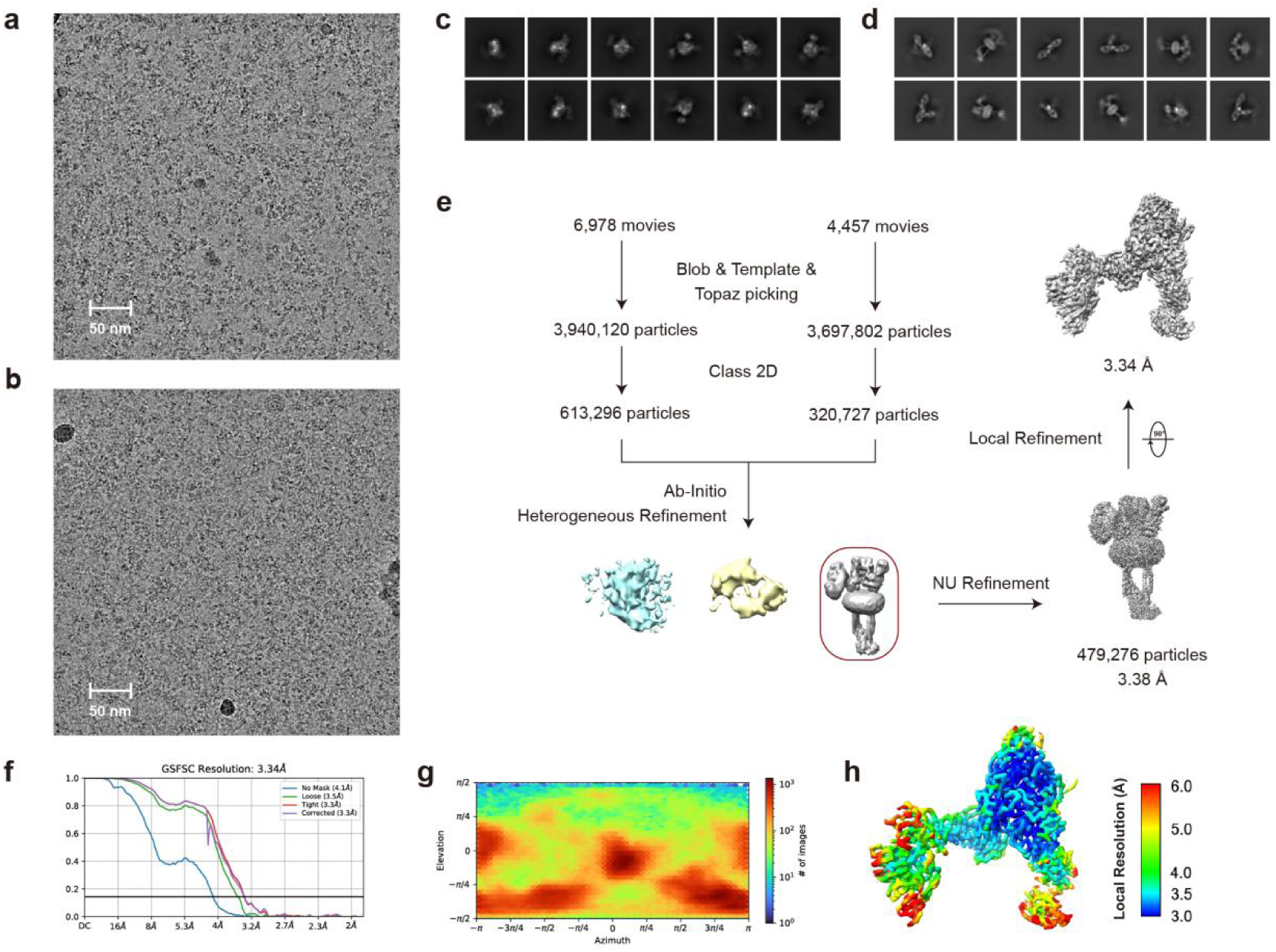
Cryo-EM analysis of the BTN3A1-BTN3A2–BTN2A1–TH001-Fab complex. **a-b.** Representative cryo-EM micrograph of the complex on holey-carbon gold grids (**a**) and GO-coated holey-carbon gold grids (**b**). Scale bar: 50 nm. **c-d.** 2D class averages of the BTN3A1-BTN3A2–BTN2A1–TH001-Fab complex on holey-carbon gold grids (**c**) and GO-coated holey-carbon gold grids (**d**). **e.** Flowchart of the cryo-EM analysis for the BTN3A1-BTN3A2–BTN2A1–TH001-Fab complex. **f.** Gold standard Fourier shell correlation (GSFSC) curves of the final density map. **g.** Angular particle distribution heat map in the final round of local refinement. **h.** Local resolution estimation of the final density map.

**Extended Data Fig. 6.**
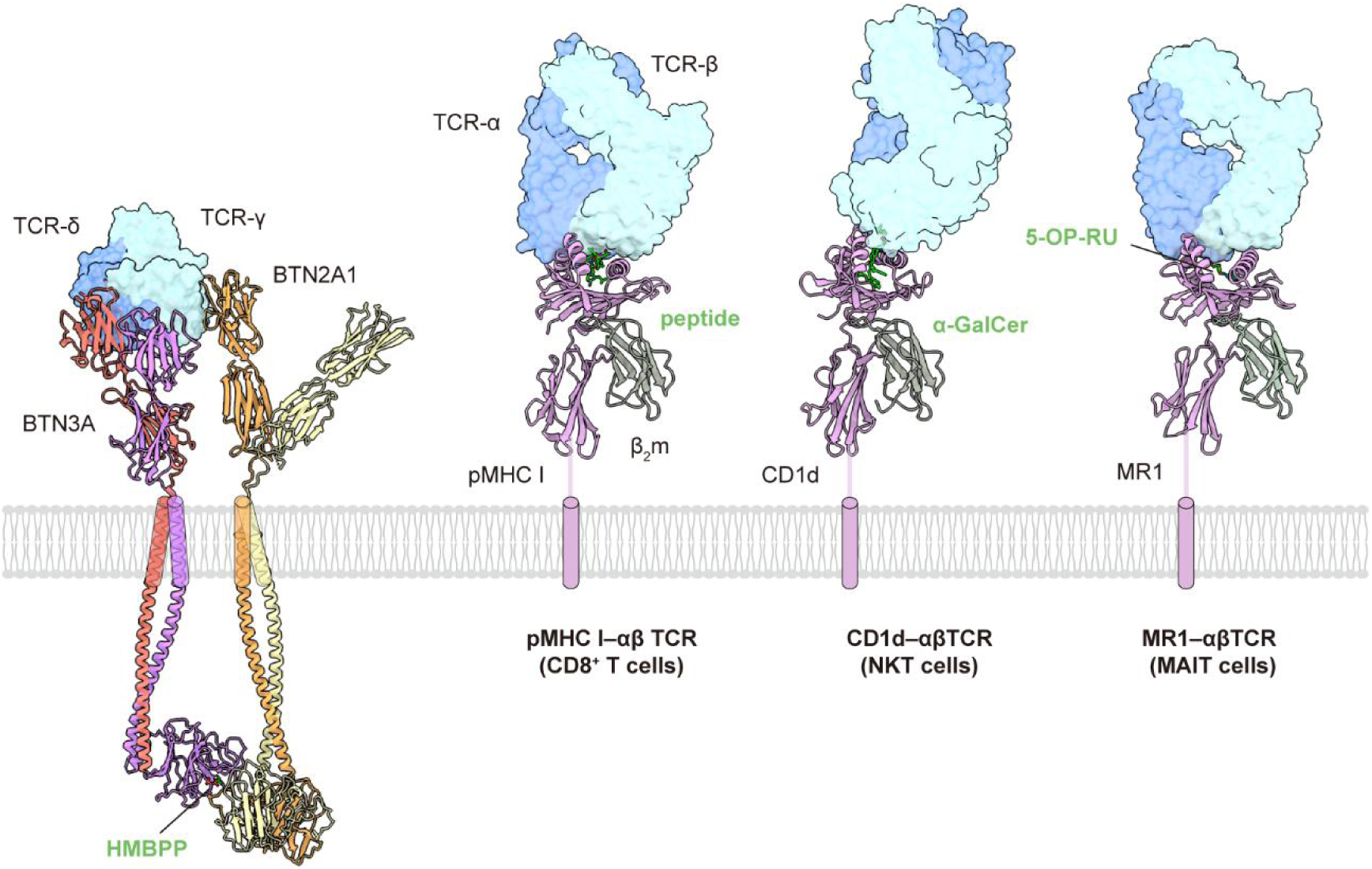
Different recognition patterns of T-cell antigens. A schematic diagram illustrating differences in TCR recognition patterns for diverse antigens, for cases including: BTN3A1-BTN3A2–HMBPP–BTN2A1–G115 Vγ9Vδ2 TCR (the present study); pMHC Ⅰ–αβ TCR (PDB ID: 7PHR)^24^; CD1d–α-GalCer–NKT αβTCR (PDB ID: 2PO6)^25^; MR1–5-OP-RU–MAIT αβTCR (PDB ID: 6PUC)^26^. Different TCR molecules are shown as surface representations in two shades of blue, while corresponding antigens on the membrane are shown as ribbons. Molecules that mediate or enhance T-cell activation are shown as green sticks.

**Extended Data Fig. 7.**
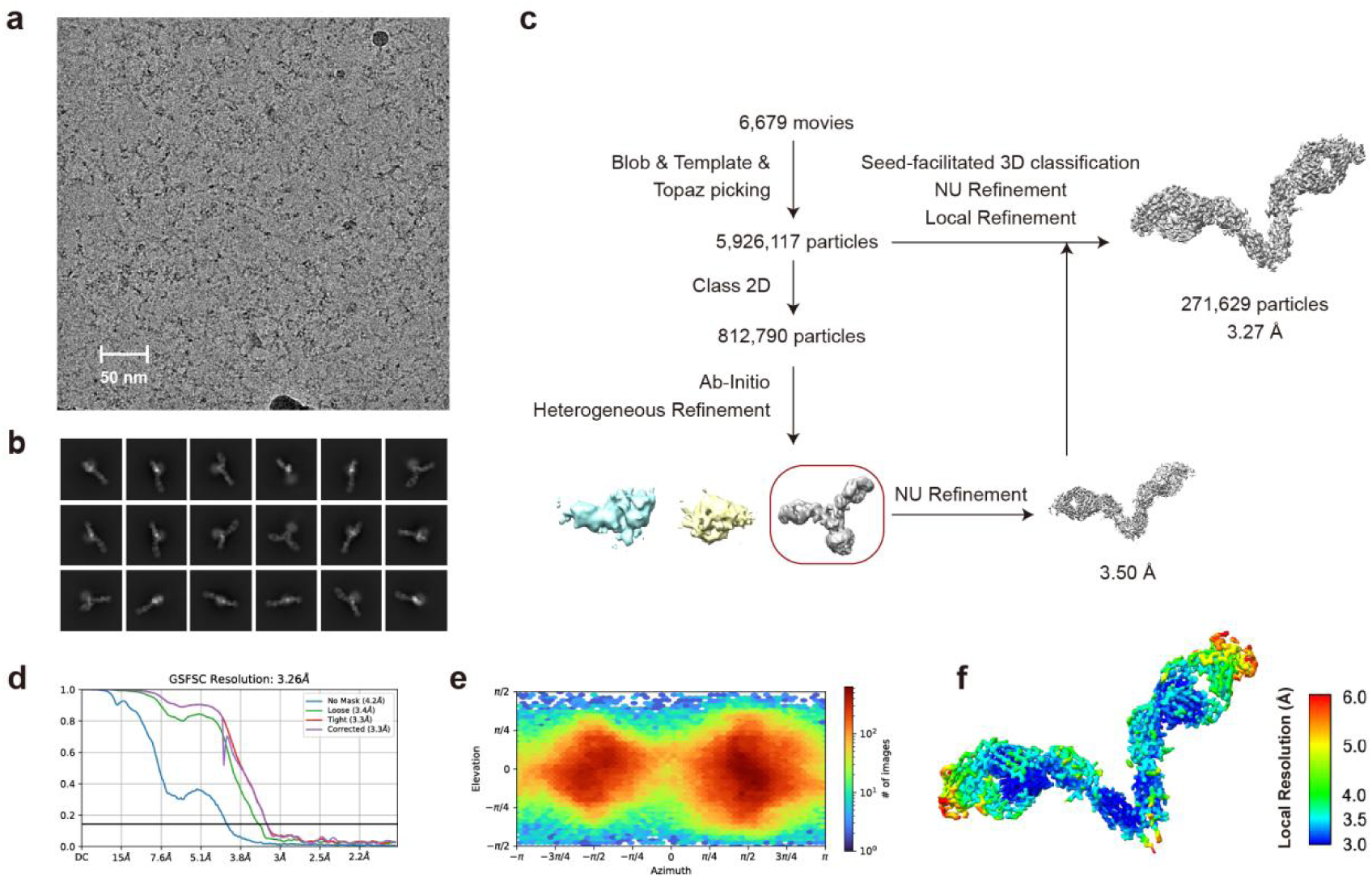
Cryo-EM analysis of the BTN2A1–TH002-Fab complex. **a-b.** Representative cryo-EM micrograph (**a**) and 2D class averages (**b**) of the BTN2A1–TH002-Fab. Scale bar: 50 nm. **c.** Flowchart of the cryo-EM analysis for BTN2A1–TH002-Fab complex. **d.** Gold standard Fourier shell correlation (GSFSC) curves of the final density map. **e.** Angular particle distribution heat map in the final round of local refinement. **f.** Local resolution estimation of the final density map.

**Extended Data Fig. 8.**
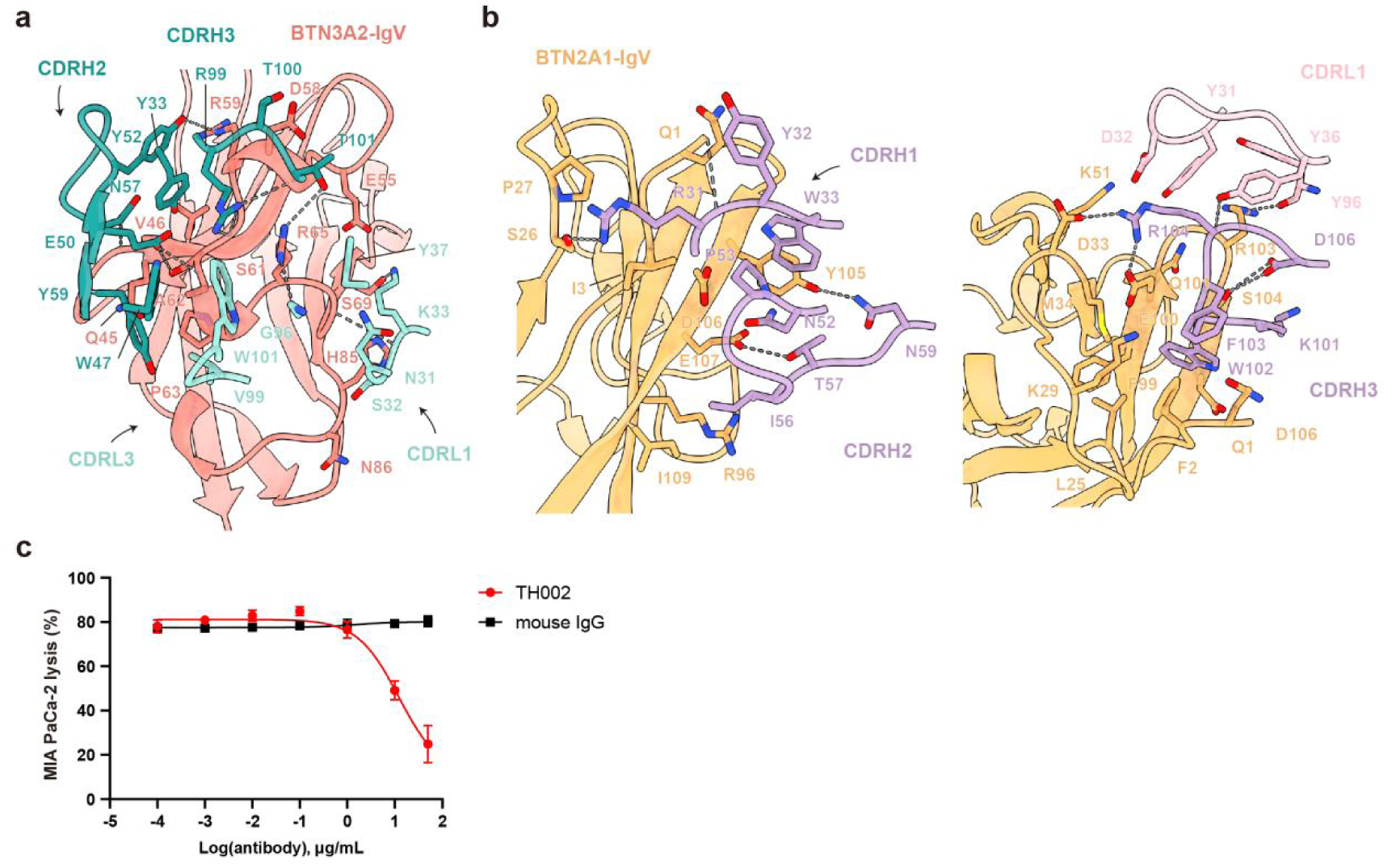
An antagonistic antibody against BTN2A1 (TH002) effectively blocks the Vγ9Vδ2 T cell activation. **a.** Interactions between TH001 and BTN3A2. Dashed lines indicate polar interactions. **b.** Interactions between TH002 and BTN2A1. **c.** TH002 blocks the responsiveness of Vγ9Vδ2 T cells towards MIA PaCa-2 cells in the presence of mAb ICT01 (1 μg/mL). Mouse IgG (IgG2a) was used as the isotype control for TH002. Representative of three independent experiments. All error bars denote the SEM.

**Extended Data Table 1.**
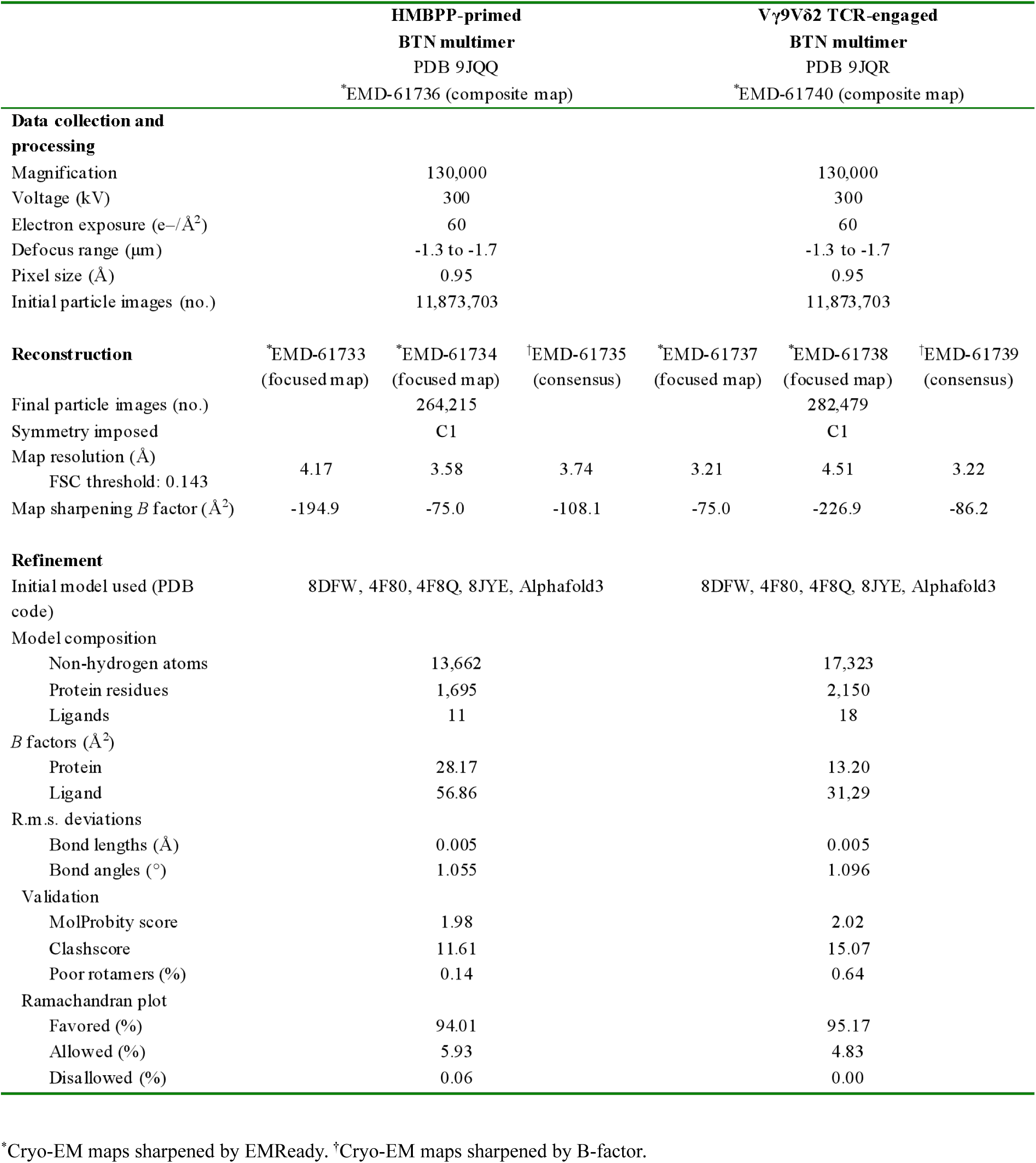
Cryo-EM analysis and validation statistics 886 of the HMBPP-primed and Vγ9Vδ2 TCR-engaged BTN3A1-BTN3A2–BTN2A1 complexes.

**Extended Data Table 2.** Cryo-EM analysis and validation statistics of the BTN3A1-BTN3A2–897 BTN2A1–TH001-Fab and BTN2A1–TH002-Fab complexes.

|  | BTN3A1-BTN3A2<br>–BTN2A1–TH001-Fab | BTN2A1–TH002-Fab |
| --- | --- | --- |
|  | PDB 9JQP<br>*EMD-61732 | PDB 9JQ6<br>†EMD-61724 |
| **Data collection and processing** |  |  |
| Magnification | 130,000 | 130,000 |
| Voltage (kV) | 300 | 300 |
| Electron exposure (e<sup>−</sup>/Å<sup>2</sup>) | 60 | 60 |
| Defocus range (μm) | -1.3 to -1.7 | -1.3 to -1.7 |
| Pixel size (Å) | 0.95 | 0.95 |
| Symmetry imposed | C1 | C1 |
| Initial particle images (no.) | 7,637,922 | 5,926,117 |
| Final particle images (no.) | 479,276 | 271,629 |
| Map resolution (Å) | 3.34 | 3.27 |
| FSC threshold 0.143 |  |  |
| **Refinement** |  |  |
| Initial model used (PDB code) | 8DFW, 4F80, 4F8Q, Alphafold3 | 8DFW, Alphafold3 |
| Model resolution (Å) | 3.34 | 3.27 |
| FSC threshold 0.143 |  |  |
| Map sharpening *B* factor (Å<sup>2</sup>) | -83.5 | -71.4 |
| Model composition |  |  |
| Non-hydrogen atoms | 8,217 | 10,277 |
| Protein residues | 1,056 | 1,304 |
| Ligands | 6 | 15 |
| *B* factors (Å<sup>2</sup>) |  |  |
| Protein | 17.42 | 12.84 |
| Ligand | 25.37 | 20.88 |
| R.m.s. deviations |  |  |
| Bond lengths (Å) | 0.005 | 0.006 |
| Bond angles (°) | 1.024 | 1.030 |
| Validation |  |  |
| MolProbity score | 2.00 | 2.03 |
| Clashscore | 11.12 | 10.76 |
| Poor rotamers (%) | 0.45 | 0.44 |
| Ramachandran plot |  |  |
| Favored (%) | 93.15 | 92.29 |
| Allowed (%) | 6.85 | 7.55 |
| Disallowed (%) | 0.00 | 0.16 |

\*Cryo-EM maps sharpened by EMReady. †Cryo-EM maps sharpened by B-factor.

